# Phylogeny and species delimitation of ciliates in the genus *Spirostomum* (Class, Heterotrichea) using single-cell transcriptomes

**DOI:** 10.1101/2024.05.29.596006

**Authors:** Shahed Uddin Ahmed Shazib, Auden Cote-L’Heureux, Ragib Ahsan, Sergio A. Muñoz-Gómez, JunMo Lee, Laura A. Katz, Mann Kyoon Shin

## Abstract

Ciliates are single-celled microbial eukaryotes that diverged from other eukaryotic lineages over a billion years ago. The extensive evolutionary timespan of ciliate has led to enormous genetic and phenotypic changes, contributing significantly to their high level of diversity. Recent analyses based on molecular data have revealed numerous cases of cryptic species complexes in different ciliate lineages, demonstrating the need for a robust approach to delimit species boundaries and elucidate phylogenetic relationships. Heterotrich ciliate species of the genus *Spirostomum* are abundant in freshwater and brackish environments and are commonly used as biological indicators for assessing water quality. However, some *Spirostomum* species are difficult to identify due to a lack of distinguishable morphological characteristics, and the existence of cryptic species in this genus remains largely unexplored. Previous phylogenetic studies have focused on only a few loci, namely the ribosomal RNA genes, alpha-tubulin, and mitochondrial CO1. In this study, we obtained single-cell transcriptome of 25 *Spirostomum* species populations (representing six morphospecies) sampled from South Korea and the USA, and used concatenation- and coalescent-based methods for species tree inference and delimitation. Phylogenomic analysis of 37 *Spirostomum* populations and 265 protein-coding genes provided a robustious insight into the evolutionary relationships among *Spirostomum* species and confirmed that species with moniliform and compact macronucleus each form a distinct monophyletic lineage. Furthermore, the multispecies coalescent (MSC) model suggests that there are at least nine cryptic species in the *Spirostomum* genus, three in *S. minus*, two in *S. ambiguum, S. subtilis*, and *S. teres* each. Overall, our fine sampling of closely related *Spirostomum* populations and wide scRNA-seq allowed us to demonstrate the hidden crypticity of species within the genus *Spirostomum*, and to resolve and provide much stronger support than hitherto to the phylogeny of this important ciliate genus.

## Introduction

Unicellular ciliates are among the most diverse clades of eukaryotes and are an essential component in both aquatic and terrestrial food webs (Hakenkamp and Morin 2000; Dopheide et al. 2008; Lynn 2008; Clamp and Lynn 2017). They are some of the most complex single-celled eukaryotes in terms of their cell structure and have even more intricate genomic architectures than multicellular eukaryotes. Ciliates of the genus *Spirostomum* Ehrenberg, 1834 (Class: Heterotrichea) are large cells (150 – 4,000 µm long) that inhabit fresh and brackish waters (Specht 1934; Nałecz-Jawecki and Sawicki 1999; Finlay 2002; Bradley et al. 2010). The genus *Spirostomum* is comprised of ten morphospecies characterized by vermiform, cylindrical and laterally flattened cell bodies, ciliary rows that spiral during contraction, and a long collecting canal that leads to a posterior contractile vacuole (Jang et al. 2012; Fernandes and Silva Neto 2013; Boscaro et al. 2014; Shazib et al. 2016, 2019; Chi et al. 2020, 2022). These “worm-like” ciliates are considered model organisms for studying ciliate ecology and physiology due to their relative abundance and sensitivity to abiotic factors (Nałecz-Jawecki and Sawicki 1999; Berger and Foissner 2003; Nałęcz-Jawecki et al. 2020). It was recently reported that many *Spirostomum* species use a rhodoquinone-dependent pathway under low oxygen conditions for anaerobic fumarate reduction (Hines et al. 2018; Mukhtar et al. 2020). *Spirostomum* species are suitable for endosymbiosis research as they can harbor either or both bacteria and eukaryotes in their cytoplasm (Fokin et al. 2005). Several *Rhizobiales* and *Rickettsiales*-like bacteria have been found in *Spirostomum* species; *S. semivirescens* is the only *Spirostomum* species reported to have *Chlorella*-like endosymbiotic green algae (Kreutz and Foissner 2006; Esteban et al. 2009; Fokin 2012; Schrallhammer et al. 2013; Hines et al. 2018; Akter et al. 2020). A unique symbiosis phylogenetically sister and morphologically similar to *Spirostomum*, *Pseudoblepahrisma tenue*, harbors contrasting photosynthetic symbionts in its cytoplasm, purple bacteria and green algae (Muñoz-Gómez et al. 2021; Muñoz-Gómez and Hess 2023). Therefore, understanding the evolution of *Spirostomum* species is also key to shedding light on the evolution of the extraordinarily rare purple-green photosymbiosis of *P. tenue*.

Several authors have previously attempted to solve the phylogenetic relationships in the genus *Spirostomum* using morphological characters and a single or a few genes (Boscaro et al. 2014; Chi et al. 2020; Fernandes and Silva Neto 2013; Shazib et al. 2019, 2016). These single and multigene analyses suggested the presence of cryptic species within the genus *Spirostomum*. For example, *S. minus* exhibits considerable diversity at the molecular level and may contain at least two morphologically undistinguishable cryptic species. *Spirostomum teres* also appears to showcase species crypticity, and has further been claimed to be a polyphyletic taxon (Boscaro et al. 2014; Shazib et al. 2019). Ribosomal RNA gene sequences may not have enough phylogenetic signal to reveal species boundaries, which were not unambiguously resolved at the molecular level using both the primary and secondary structures of rRNA gene sequences (Shazib et al. 2016). In our recent work, we increased the taxon sampling and used two protein-coding genes, i.e., nuclear alpha-tubulin and mitochondrial CO1, to delimit *Spirostomum* species (Shazib et al. 2019). We also used the Bayesian coalescent approach to estimate a species tree, but neither nuclear nor mitochondrial genes could definitively resolve the inter-relationships among *Spirostomum* species due to insufficient phylogenetic signal. The multispecies coalescent (MSC) model provides a powerful approach to delimit species based on molecular data. Thus, additional genomic information and analytical approaches are required to understand the species complex structure and resolve the evolutionary relationships of the *Spirostomum* genus.

Here, we generated 25 new single-cell transcriptomes of *Spirostomum* populations from fresh and brackish water environments in South Korea and the USA. We used these, together with 12 additional publicly available transcriptomes, belonging to seven morphospecies (*S. ambiguum, S. caudatum, S. minus, S. teres, S. semivirescens, S. subtilis, S. yagiui*), to infer a robust phylogenetic tree and investigate the species boundaries of the genus *Spirostomum*. To do this, we thus performed concatenation- and coalescent-based analyses on a dataset that encompasses 37 *Spirostomum* populations and 265 protein-coding genes. We also assembled a pipeline to reconstruct the rRNA gene sequences from transcriptomic data, as the rRNA gene is key to understanding species diversity more broadly. Our study represents the first attempt to resolve the phylogeny and species boundaries of the *Spirotsomum* genus using single-cell transcriptomes.

## Materials and Methods

### Sample collection, isolation, and culturing

Samples were collected from various sites, including fresh and brackish waters (Table 1). After transportation to the laboratory, the collected samples were immediately stored in a cool box and examined directly under a dissecting microscope. Single cells of each ciliate morphospecies were isolated, washed in sterile distilled water, and used to start new cultures. The cultures were maintained at 20-24 °C in Petri dishes, and wheat grains were periodically added to stimulate the growth of prey bacteria. Culture dishes contained filtered *in-situ* water or commercial mineral water (Evian, France) for freshwater species, and sterile seawater for marine/brackish water species. Specimens from each clonal culture were examined under a light microscope, using bright-field and differential interference contrast (DIC) optics. Species characterization and identification were performed according to the following guidelines: Repak and Isquith (1974); Foissner et al. (1992); Berger et al. (1997); Boscaro et al. (2014); Shazib et al. (2019); Chi et al. (2020) (Supplementary Fig. S1).

**Table 1.** Summary of *Spirostomum* species and transcriptomes included in the present work.

| MAC pattern | Species | Population ID | sc RNA-seq protocol | Raw reads (M) | Eukaryotic Contigs* | BUSCO Analysis (%) | GenBank accession # | Country | Habitat | Reference |
| --- | --- | --- | --- | --- | --- | --- | --- | --- | --- | --- |
| Moniliform | *S. ambiguum* | Sa_C1 | Total RNA extraction | 61 | 16,523 | 87.7 | SRR12125180 | China | Fresh water | Mukhtar et al., 2020 |
| Moniliform | *S. ambiguum* | Sa_U1 | SMART-Seq v4 | 48 | 9,567 | 90.7 | SRR10512970 | USA | Fresh water | Yan et al., 2019 |
| Moniliform | *S. ambiguum* | Sa_K1 | SMART-Seq v4 | 48 | 7,421 | 80.1 | XXXXXXXXXX | South Korea | Fresh water | This study |
| Moniliform | *S. ambiguum* | Sa_K2 | SMART-Seq v4 | 53 | 8,531 | 86.5 | XXXXXXXXXX | South Korea | Fresh water | This study |
| Moniliform | *S. ambiguum* | Sa_K3 | SMART-Seq v4 | 106 | 8,510 | 88.9 | XXXXXXXXXX | South Korea | Fresh water | This study |
| Moniliform | *S. ambiguum* | Sa_K4 | Smart-seq2 | 128 | 9,515 | 88.3 | XXXXXXXXXX | South Korea | Fresh water | This study |
| Moniliform | *S. ambiguum* | Sa_K5 | Smart-seq2 | 124 | 7,070 | 81.3 | XXXXXXXXXX | South Korea | Fresh water | This study |
| Moniliform | *S. ambiguum* | Sa_K6 | Smart-seq2 | 114 | 8,140 | 87.2 | XXXXXXXXXX | South Korea | Fresh water | This study |
| Moniliform | *S. ambiguum* | Sa_K7 | Smart-seq2 | 48 | 8,580 | 84.1 | XXXXXXXXXX | South Korea | Fresh water | This study |
| Ellipsoid | *S. caudatum* | Sc_K1 | Smart-seq2 | 70 | 2,078 | 29.3 | XXXXXXXXXX | South Korea | Brackish water | This study |
| Moniliform | *S. minus* | Sm_U1 | SMART-Seq v4 | 59 | 3,789 | 63.7 | SRR10513010 | USA | Fresh water | Yan et al., 2019 |
| Moniliform | *S. minus* | Sm_U6 | SMART-Seq v4 | 13 | 7,085 | 82.5 | XXXXXXXXXX | USA | Fresh water | This study |
| Moniliform | *S. minus* | Sm_U25 | SMART-Seq v4 | 36 | 2,091 | 45.1 | XXXXXXXXXX | USA | Fresh water | This study |
| Moniliform | *S. minus* | Sm_K1 | Smart-seq2 | 54 | 4,969 | 72.6 | XXXXXXXXXX | South Korea | Fresh water | This study |
| Moniliform | *S. cf. minus* | Sm_K2 | Smart-seq2 | 73 | 11,040 | 93 | XXXXXXXXXX | South Korea | Fresh water | This study |
| Moniliform | *S. minus* | Sm_K3 | Smart-seq2 | 71 | 8,456 | 83 | XXXXXXXXXX | South Korea | Fresh water | This study |
| Moniliform | *S. minus* | Sm_K4 | Smart-seq2 | 85 | 10,188 | 88.3 | XXXXXXXXXX | South Korea | Fresh water | This study |
| Moniliform | *S. sp.* | Ss_E1 | Smart-seq2 | 12 | 7,799 | 85.5 | SRR7141210 | Sweden | Fresh water | Hines et al., 2018 |
| Moniliform | *S. semivirescens* | Sv_E1 | Smart-seq2 | 2 | 2,743 | 55.6 | SRR7141204 | England | Fresh water | Hines et al., 2018 |
| Moniliform | *S. semivirescens* | Sv_E2 | Smart-seq2 | 4 | 6,454 | 77.8 | SRR7141205 | England | Fresh water | Hines et al., 2018 |
| Moniliform | *S. semivirescens* | Sv_E3 | Smart-seq2 | 4 | 5,473 | 74.9 | SRR7141206 | Sweden | Fresh water | Hines et al., 2018 |
| Moniliform | *S. semivirescens* | Sv_E4 | Smart-seq2 | 3 | 5,102 | 76 | SRR7141207 | Sweden | Fresh water | Hines et al., 2018 |
| Moniliform | *S. semivirescens* | Sv_E5 | Smart-seq2 | 1 | 3,052 | 62 | SRR7141208 | Sweden | Fresh water | Hines et al., 2018 |
| Moniliform | *S. semivirescens* | Sv_E6 | Smart-seq2 | 3 | 7,285 | 84.7 | SRR7141209 | Sweden | Fresh water | Hines et al., 2018 |
| Moniliform | *S. subtilis* | Su_U10 | SMART-Seq v4 | 12 | 9,684 | 91.8 | XXXXXXXXXX | USA | Fresh water | This study |
| Moniliform | *S. subtilis* | Su_C1 | Total RNA extraction | 51 | 17,617 | 91.2 | SRR12125179 | China | Fresh water | Mukhtar et al., 2020 |
| Moniliform | *S. subtilis* | Su_K1 | Smart-seq2 | 56 | 9,110 | 86 | XXXXXXXXXX | South Korea | Fresh water | This study |
| Moniliform | *S. subtilis* | Su_K2 | Smart-seq2 | 80 | 8,591 | 85.4 | XXXXXXXXXX | South Korea | Fresh water | This study |
| Moniliform | *S. subtilis* | Su_K3 | Smart-seq2 | 51 | 10,216 | 93 | XXXXXXXXXX | South Korea | Fresh water | This study |
| Ellipsoid | *S. teres* | St_C1 | Total RNA extraction | 53 | 14,595 | 94.2 | SRR12125178 | China | Fresh water | Mukhtar et al., 2020 |
| Ellipsoid | *S. teres* | St_K1 | Smart-seq2 | 54 | 5,336 | 72.5 | XXXXXXXXXX | South Korea | Fresh water | This study |
| Ellipsoid | *S. teres* | St_K3 | SMART-Seq v4 | 53 | 7,929 | 84.9 | XXXXXXXXXX | South Korea | Fresh water | This study |
| Elongated | *S. yagiui* | Sy_K1 | Smart-seq2 | 54 | 6,037 | 78.4 | XXXXXXXXXX | South Korea | Fresh water | This study |
| Elongated | *S. yagiui* | Sy_K2 | SMART-Seq v4 | 58 | 6,396 | 80.7 | XXXXXXXXXX | South Korea | Fresh water | This study |
| Elongated | *S. yagiui* | Sy_K3 | Smart-seq2 | 165 | 7,594 | 84.8 | XXXXXXXXXX | South Korea | Fresh water | This study |
| Elongated | *S. yagiui* | Sy_K4 | Smart-seq2 | 109 | 7,558 | 81.3 | XXXXXXXXXX | South Korea | Fresh water | This study |
| Elongated | *S. yagiui* | Sy_K5 | Smart-seq2 | 113 | 7,118 | 87.8 | XXXXXXXXXX | South Korea | Fresh water | This study |
| Moniliform | *Anigstenia* sp. | As_U1 | SMART-Seq v4 | 183 | 2,877 | 33.9 | SRR10512978 | USA | Marine water | Yan et al., 2019 |

M = million.; Read Mapping (RM) = rDNA reconstructed from transcriptomic data; BUSCO analysis = Completeness + Partial, \* = number of eukaryotic contigs after PhyloToL pipeline part 1 (PLT1) (see methods and materials).

### Single-cell preparation

Single cells were isolated with a capillary glass pipette under a dissecting microscope. Cells were washed three times in sterile distilled water through successive transfers in PYREX 6-well depression plates, diluting the original medium and removing eukaryotic and bacterial contaminants. Ciliates from brackish habitats were washed three times in sterile *in situ* water. In the last step, cells were washed with distilled water and transferred to 1.5 mL tubes for DNA extraction and 0.2 mL tubes for single-cell transcriptome amplification. Cells were frozen at -80°C or processed immediately.

### Single-cell DNA extraction

DNA was extracted using the RED Extract-N-Amp Tissue PCR Kit (Sigma, St. Louis, MO) according to the protocol of Shazib et al. (2019). The rRNA genes (18S, ITS, 28S) were amplified by polymerase chain reaction (PCR) using the TaKaRa Ex Taq polymerase kit (TaKaRa Bio-medicals, Seoul, Korea). PCR cycling parameters and primer information were followed as in Shazib et al. (2019). PCR products were directly sequenced on an ABI 3730 automated sequencer (Macrogen Inc., Seoul, South Korea) using seven sequencing primers (Supplementary Table S1). Sequence fragments from individual samples were checked, trimmed, and assembled into contigs using Geneious ver. Prime 2019 (Biomatters, http://www.geneious.com/) (Kearse et al. 2012).

### Single-cell transcriptomes

Single-cell Whole Transcriptome Amplifications (scWTA) were performed from frozen cells (-80°C) or cells freshly picked from the clonal cultures. WTAs of the individual ciliates were generated according to the Smart-seq2 protocol (Picelli et al. 2014) and SMART-Seq v4 Ultra Low Input RNA Kit (TaKaRa Bio-medicals, Seoul, Korea) using 17 cycles of cDNA amplification for each cell. After transcriptome amplification, cDNA was further cleaned using the ProNex® Size-Selective Purification System (Promega, WI, USA) according to the manufacturer’s protocol. Following ProNex® purification, samples were quantified with a Qubit 2.0 Fluorometer (ThermoFisher Scientific, USA). Sequencing libraries were constructed using a Nextera XT DNA library preparation kit (Illumina, FC-131-1024) and a Nextera XT Index kit (Illumina, FC-131-1001). Sequencing was done on a HiSeq 2500 (100 bp PE) and NovaSeq 6000 (150 bp PE) instruments at Theragen Bio (Seoul, South Korea).

Whole transcriptome amplification for three *Spirostomum* populations from the Katz lab was performed using the SMART-Seq® v4 Ultra® Low Input RNA Kit for Sequencing (TaKaRa Bio USA, Inc., Mountain View, CA, USA). Amplified cDNA was purified using the AMPure XP PCR Purification system (Beckman Coulter Life Sciences, Indianapolis, IN, USA) and then quantified using a Qubit 3.0 Fluorometer (ThermoFisher Scientific, Waltham, MA, USA). The Nextera XT DNA library preparation kit (Illumina, FC-131-1096) and Nextera XT Index kit v2 set A (Illumina, FC-131-2001) were used to construct Illumina sequencing libraries. These were then sequenced on a HiSeq 4000 instrument (Illumina) at the Institute for Genome Sciences, University of Maryland (Baltimore, USA).

### High-Throughput Sequencing data processing

We assessed the quality of sequenced raw reads using FastQC v.0.11.5 (Andrews 2010), and trimmed adapters and removed low-quality sequences with BBtools (Bushnell et al. 2017) using q24 as minimum quality and minimum length (125bp for 150bp, 300bp PE reads and 70 bp for 100bp PE reads). Transcriptome assembly was carried out using rnaSPAdes (Bushmanova et al. 2019) and subsequently processed by a series of custom Python scripts from PhyloToL (PhyloToL part 1-Post assembly Pipeline) (Cerón-Romero et al. 2019), available at https://github.com/Katzlab. The post assembly pipeline removes contaminants (rDNA and non-eukaryotic sequences) and translates the nucleotide sequences with >200bp after assigning amino acids to GFs (gene families) with an appropriate genetic code. Further, we assessed the completeness of the translated sequences using BUSCO ver. 5.4.7 (Alveolata dataset −l alveolata_odb10, and proteome mode -m protein) (Manni et al. 2021) (Table 1; Supplementary Table S3).

### rDNA reconstruction, taxonomy assignment, and tree construction

We sequenced the rRNA gene operon (18S, ITS, 28S) of 21 populations using the Sanger method (Supplementary Table S5). In addition, we reconstructed the rRNA gene sequences from other *Spirostomum* and *Anigsteinia* populations by mapping reads from their respective raw transcriptomes against a set of reference sequences downloaded from NCBI (87 reliable rRNA gene sequences of the genus *Spirostomum* and *Anigsteinia* (Shazib et al. 2019). We tested various mapping software, that include Bowtie 2 (Langmead and Salzberg 2012), BBMap (Bushnell 2014), BWA (Li and Durbin 2009), and HISAT2 (Kim et al. 2019), to assess their influence on the accuracy of the reconstructed contigs. After read mapping to the 18S rRNA gene, three different read sets with differing quality thresholds (all reads, q20, and q40) were used for read assembly with rnaSPAdes (Bankevich et al. 2012; Bushmanova et al. 2019) (Supplementary Table S4). We aligned all contigs reconstructed using various read mappers and reference sequences to identify errors and artifacts (Supplementary Fig. S3). To construct rRNA gene sequence data matrix, all reliable *Spirostomum* sequences aligned with newly obtained sequences using the MUSCLE algorithm in Geneious ver. Prime 2019 (Edgar 2004). Ambiguous sequences and poorly aligned columns were removed from the data matrix generating a final multiple sequence alignment (MSA) of 2,459 bp which was then analyzed with IQ-TREE ver. 2.2.0.3 to construct a maximum likelihood (ML) tree (Minh et al. 2020). The TIM2e+I+G4 model was the best substitution model based on the Bayesian information criterion (BIC) calculated in IQ-TREE ver. 2.2.0.3. Maximum likelihood (ML) analyses were carried out using 1,000 replicates in IQ-TREE. Phylogenetic trees were visualized and edited using FigTree ver. 1.4 (http://tree.bio.ed.ac.uk/software/figtree/). The reliability of branch support was assessed using ultrafast bootstrap approximation (UFBoot), and the Shimodaira Hasegawa-like approximate likelihood ratio test (SH-aLRT) with 1,000 replicates in both methods (Guindon et al. 2010; Minh et al. 2013). The reliability of a clade only be considered if the Ufboot support and SH-aLRT values are ≥ 95% and ≥ 80%, respectively (Guindon et al. 2010; Minh et al. 2013).

### Phylogenomic dataset construction

All publicly available *Spirostomum* transcriptomes were downloaded from NCBI GenBank as of April 2021, processed, and added to the PhyloToL database (Table 1; Supplementary Table S2). Initially, we generated 1,245 gene trees using PhyloToL and assessed transcript diversity in each family for each species. Bacterial and eukaryotic contamination was identified by analyzing single-gene trees summarized by a custom Python script (ContaminationBySisters.py). Next, we inspected paralogous families in these initial single-gene trees and picked out the 1,141 best clades (subtrees) with only target sequences using the custom Python script (GetBestSequences.py). Sequences with short lengths and low coverages (cov ≤ 10) were removed before alignments and trees were reconstructed. After manual hand curation, we selected 265 subtrees with sequences that include single-copy genes for each species (Gene tree inspection and curation steps in Fig. 1; Supplementary Table S8). The PhyloToL scripts used above are available at https://github.com/Katzlab/PhyloToL-6/tree/main/Utilities.

**Figure 1:**
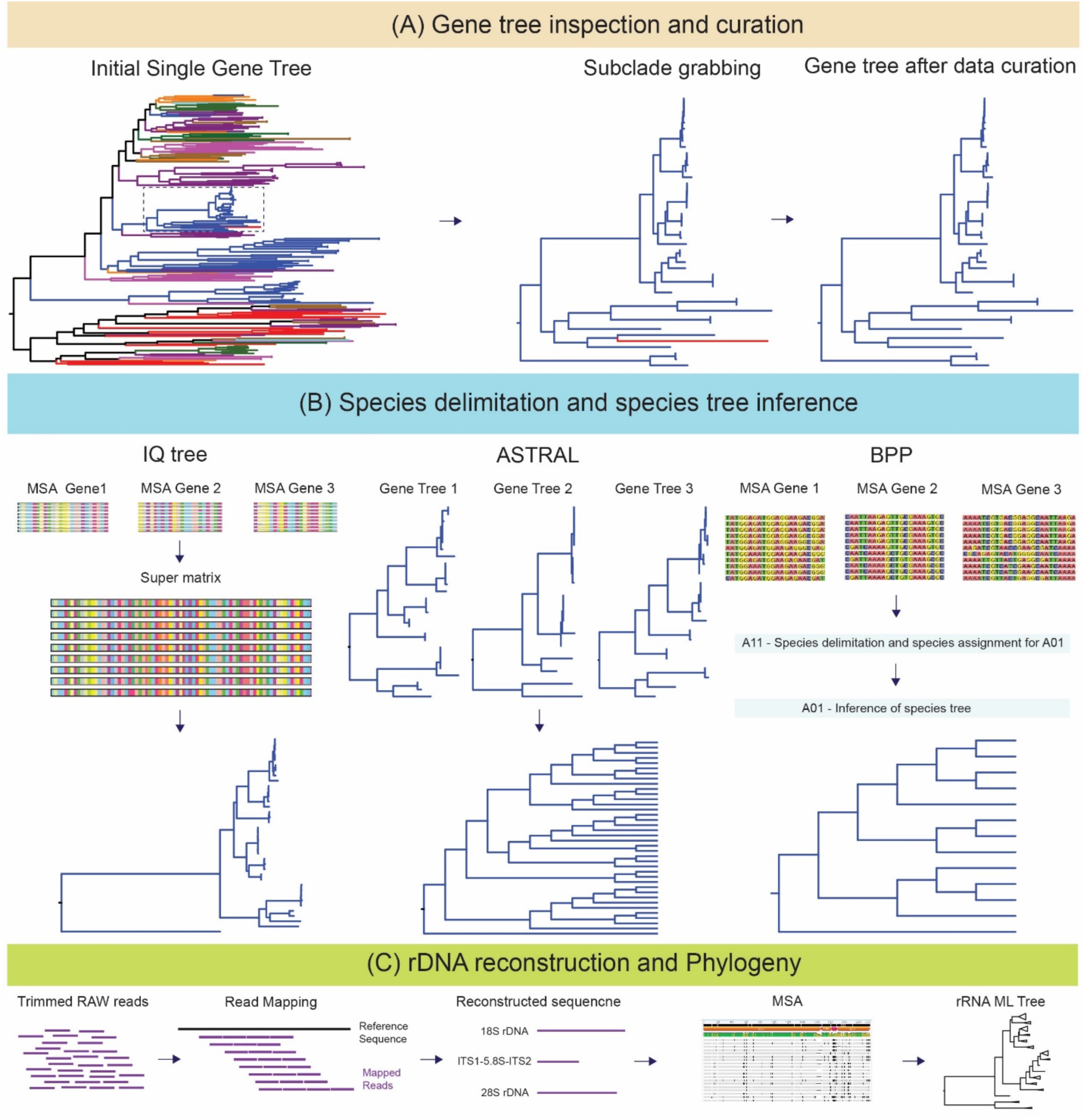
Schematic workflow for phylogenomic analysis and reconstruction of rRNA genes from single-cell transcriptomic data. (A) Initial gene trees were constructed using the taxon/gene-rich pipeline PhyloToL. Each tree was inspected, and subclades containing maximum numbers of target taxa were extracted. Subclades were manually checked for non-target taxa. Curated gene trees with single-copy gene from all taxa were used to prepare the data matrix. (B) For concatenation analysis, each dataset with well-curated sequences were aligned and 265 datasets were concatenated into an amino acid super matrix for ML phylogeny. The ASTRAL method used 265 protein-coding gene trees to infer a species tree. Species delimitation and species tree inference were performed using BPP with 265 well-aligned datasets of nucleotides (C) rRNA gene sequences were reconstructed from raw single-cell RNA sequencing (scRNA-seq) data using read mapping software (Hisat2 and Bowtie 2). Reconstructed sequences were aligned, and rRNA phylogeny was inferred using the ML approach. Detailed methods are provided in the Methods and Materials section.

### Concatenation-based phylogenomic approach

The well-curated datasets of 265 orthologous genes from 38 taxa were aligned with MAFFT using the L-INS-I option (Katoh and Standley 2013). All MSAs were inspected, and gaps were removed using trimAl (-gt 0.9) (Capella-Gutiérrez et al. 2009). Concatenation of 265 MSAs into a super matrix were performed using Geneious ver. Prime 2019 (Kearse et al. 2012). The final dataset comprises 62,043 unambiguously aligned amino acid residues. Running a phylogenetic analysis using the site-heterogeneous CAT-GTR model in PhyloBayes is often computationally demanding for a 265 protein-coding concatenated dataset. Therefore, only maximum likelihood (ML) analyses were performed for the large dataset. The ML phylogenetic analyses were performed using IQ-TREE ver. 2.2.0.3 using the LG +C60 + F + G4 profile mixture model (Si Quang et al. 2008). To estimate of branch support, we used the ultrafast bootstrap approximation (UFBoot) with 1,000 replicates (Minh et al. 2013; Hoang et al. 2018) and the SH-like approximate likelihood ratio test (SH-aLRT) also with 1,000 bootstraps (Guindon et al. 2010).

To visualize potential phylogenetic conflicts or noise in the concatenated 265 protein-coding gene dataset, a neighbor-net network analysis was performed with uncorrected distances in SplitsTree CE (Huson 1998; Huson and Bryant 2006). The reliability of splits was assessed with 1,000 bootstrap replicates (Bryant 2003).

### Coalescent-based species delimitation and species tree analyses

Discordance between gene and species tree occurs due to several biological factors such as incomplete lineage sorting (ILS) and introgression. Such discordance can be minimized by using coalescent-based methods (Huang et al. 2020). We used two different approaches to estimate the species tree using the multispecies coalescent model (MSC). First, we used the Accurate Species TRee Algorithm (ASTRAL) III ver. 5.7.1 software which uses a multispecies coalescent (MSC) model to account the incomplete lineage sorting (ILS) (Zhang et al. 2018) and infer species tree from the protein-coding gene trees. This method is widely used, and the accuracy of the species tree relies on the given error-free input gene trees (Mirarab et al. 2014). We constructed individual gene trees from 265 protein-coding gene alignments using ML (maximum likelihood) approach in IQ-TREE ver. 2.2.0.3. These single-gene trees were used by ASTRAL to estimate a species tree, and the certainty of the tree nodes were quantified using local posterior probability (LPP) (Sayyari and Mirarab 2016).

Second, we used the Bayesian Phylogenetics and Phylogeography (BPP) ver. 4.6.2 software (Yang 2015; Flouri et al. 2018) which uses a multispecies coalescent (MSC) model under the Bayesian framework and estimates the posterior probabilities (PP) from the alternative species tree and a species delimitation hypothesis (Flouri et al. 2018, 2020a). Using the reversible-jump Markov chain Monte Carlo (rjMCMC) algorithm and a set of priors, BPP can combine information at many loci and estimate a reliable species tree even if every locus has a weak phylogenetic signal (Xu and Yang 2016; Shi and Yang 2018).

For species delimitation, we performed the A11 analysis (species delimitation = 1 and species tree = 1) in BPP, which accounts for joint species delimitation and species-tree estimation. The software uses nucleotide sequences, as synonymous codon changes in the nucleotide sequences provide more information when working with closely related species (Flouri et al. 2020b). Therefore, we used MSAs of DNA sequences for BPP analyses and each alignment file used as single loci input files for BPP.

The concatenated phylogenomic tree from 265 protein-coding genes was used as a guide tree for species delimitation (A11). However, estimating species tree and species delimitation should not be affected by starting tree if there is no convergence and mixing problems (Robert and Casella 2004; Yang and Rannala 2014). Large datasets like the 265 loci one used here may pose problems to MCMC convergence and mixing and thus produce inconsistent results (Yang 2015; Flouri et al. 2018; Thawornwattana et al. 2022) (Supplementary Table S8). To get the consistency and proper mixing of the across different analyses initially, we used a subset dataset of 100 loci, each of which has more than 70% of taxon occupancy, and the nucleotide size ranges between 200 bp – 1,000 bp after removing any sites with missing data (Supplementary Table S9). Using the 100 loci dataset, we set six sets of priors to explore the sensitivity of BPP analysis using α = 3 in the inverse-gamma as a diffuse prior and α = 21 as an informative prior (Flouri et al. 2018) (Supplementary Table S7). After initial assessment using six different prior settings, we assigned priors θ ∼ InvG (3, 0.002), with mean 0.001, and τ ∼ InvG (3, 0.4), with mean 0.2. We used rjMCMC analyses and algorithm 0 and the fine-tuning parameter was set as e=10 to guarantee a good mixing in the reverse jump (rj) algorithm. The rjMCMC was set to 100,000 generations with burnin = 10,000 and sample frequency 5. Posterior probabilities ≥ 0.95 were considered highly supported. Each analysis was run at least two times and convergence was assessed by examining consistency between runs. We also ran BPP for 265 loci using the same parameters for 100 loci. Both datasets will generate consistent results if the MCMC chains are properly mixed.

We performed A01 analysis (species delimitation = 0 and species tree = 1) with BPP (Yang and Rannala 2014; Yang 2015) for species phylogeny and partitioned all 38 individuals into 13 “species’’ following the results of species delimitation analyses from A11. Species tree inferences using A01 method under the MSC model have higher efficiency in estimating correct species tree than concatenation and summary-based methods when large data sets are used (Flouri et al. 2018). We used the uniform rooted tree prior (speciesmodelprior = 1) and excluded any ambiguous sites (cleandata = 1). The ML tree from the concatenated 265-gene dataset used as a guide tree for this analysis and the same prior settings used for species delimitation (A11). To check the reliability of our result, we ran each analysis at least twice. Each analysis was run for 100,000 generations, sampling every 5th generation with 10,000 burn-in generations. Posterior probabilities greater than 0.95 were considered highly supported lineage.

We used DensiTree (Bouckaert 2010) to visualize the agreements in the trees generated during the MCMC run of the A01 BPP analysis. All trees generated in the A01 analysis (200,000 MCMC trees, from two runs) were exported as Newick format files and visualized in DensiTree.

## Results

### Ciliate transcriptome amplification using scRNA-seq

We used two different methods to amplify the transcriptomes from single ciliate cells SMART-Seq V4 and Smart-seq2 (see methods). Both methods rely on polyadenylated mRNA amplification, and have previously been successfully employed on single-cell eukaryotes (Kolisko et al. 2014; Liu et al. 2017; Onsbring et al. 2020). We amplified transcriptomes from eight populations using the SMART-Seq V4 kit and 17 populations using the Smart-seq2 protocol (Table 1). Sequencing results from 35 populations using the Smart-seq technique show that both kits performed similarly, and data on contig number plus BUSCO analyses vary across populations and sequencing depth (Table 1; Supplementary Table S3).

### rDNA construction from HTS data

We reconstructed rRNA genes by recruiting reads that map against references and then reconstructing a sequence for each cell (Fig. 1; Supplementary Table S6). We found that contigs generated using BWA from *Spirostomum subtilis* (Su_K1) WTA data had poor quality, with over 50% sequence mismatch nucleotide against the reference sequences. HISAT2 produced the longest contig, with over 99% matching the reference sequences; Bowtie 2 and BBMap generated the second and third longest contigs respectively (Supplementary Fig. S3). HISAT2 and Bowtie 2 showed comparable results (only two nucleotide mismatches, and over 99.9% similarity) with the references. But BBMap showed some mismatched or erroneous sequences in the assembled contig. Meanwhile, q40 retrieved the most reliable reads except BWA, and reads from the other three mappers (BBMap, Bowtie 2 and HISAT2) generated reliable and single assembled contig (Supplementary Table S4). Our results indicate that HISAT2 and Bowtie 2, with a quality score of 40, perform the best for constructing rRNA gene from ciliate WTA data.

### rRAN gene phylogenetic tree

In the rRNA gene tree, *Spirostomum* was divided into two major clades that corresponds with the distribution of macronuclear morphology (Fig. 2; Supplementary Fig. S2). The rRNA gene tree generally supported the monophyly of all *Spirostomum* morphospecies except *S. minus* and *S. teres*. However, most of the phylogenetic relationships among *Spirostomum* morphospecies were either poor or moderately supported.

**Figure 2.**
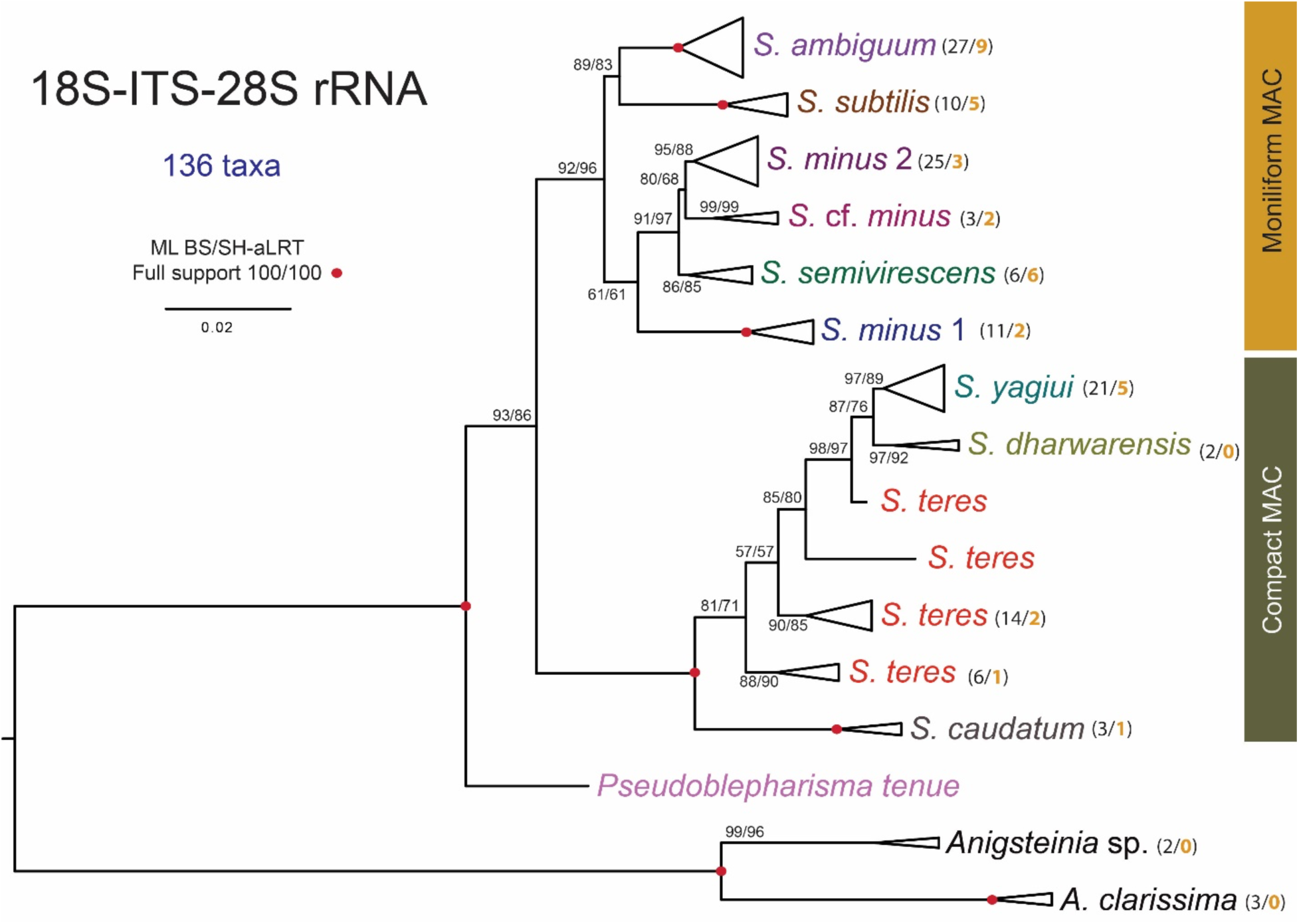
Maximum likelihood (ML) phylogenetic tree of genus *Spirostomum*. The topology is based on 136 18S-ITS-28S rRNA gene operon sequences (2,459 bp) and the TIM2e+I+G4 model. Monophyletic clades were collapsed for clarity (see also Supplementary Material Fig. S2). Numbers within brackets indicate the total number of populations (in black font) and the number of populations used for phylogenomic analyses (in bold orange font). Numbers at nodes are ML bootstrap values and SH-aLRT from 1,000 replicates. Red solid circle on branches denotes maximal support. The scale bar corresponds to the number of nucleotide substitutions.

The first clade, all sharing a moniliform macronucleus (MAC), united *S. minus*, *S. ambiguum,* and *S. subtilis* with moderate support (92% UFBoot and 96% SH-aLTR). *Spirostomum minus* splits into two clades, and the monophyly of *S. minus* clade 1 has full support, whereas *S. minus* clade 2 forms a sister relationship with *Spirostomum* cf. *minus* clade with poor statistical support (80% UFBoot and 68% SH-aLTR). *Spirostomum* cf. *minus* clade comprises three taxa (SKS255, Sm_K2, Ss_E1) and forms a highly supported monophyletic clade. *Spirostomum* cf*. minus* Sm_K2 which morphologically resemble *S. minus*, and rRNA gene tree suggest, all three taxa are close to *S. minus* clade 2 (Fig. 2; Supplementary Fig. S2). *S. subtilis* forms a clade sister to *S. ambiguum* (89% UFBoot and 83% SH-aLTR), and *S. semivirescens* falls sister to *S. minus* 2 + *Spirostomum* cf. *minus* clade with moderate supports (91% UFBoot and 97% SH-aLTR).

The second clade has lineages with compact MAC and contained *S. teres*, *S. yagiui*, *S. dharwarensis,* and *S. caudatum*, and monophyletic origin of compact macronucleus species was fully supported (Fig. 2). *S. teres* appeared to be paraphyletic, as *S. caudatum* early diverged taxa in the second clade with full statistical supports. Besides that, *Pseudoblepharisma tenue* (Pt_MG) with a compact MAC appears to have branched off first among all *Spirostomum* species, although the topology is supported by only moderate value (93% UFBoot and 86% SH-aLTR). All *Anigstenia* species formed a fully supported monophyletic clade, consistent with the classification as the sister to all *Spirostomum* spp. in all our previous phylogenetic analyses (Shazib et al. 2014; Chen et al. 2017, 2018).

### Concatenated analyses

Maximum likelihood (ML) analyses of the concatenated super matrix from 265 protein-coding genes yielded a topology with strong support for most branches (Fig. 3a). The majority of the presently defined morphospecies were monophyletic, except for *S. minus.* Both major clades identified in the rRNA gene trees were fully supported (cp. Fig. 2 and Fig. 3a).

**Figure 3.**
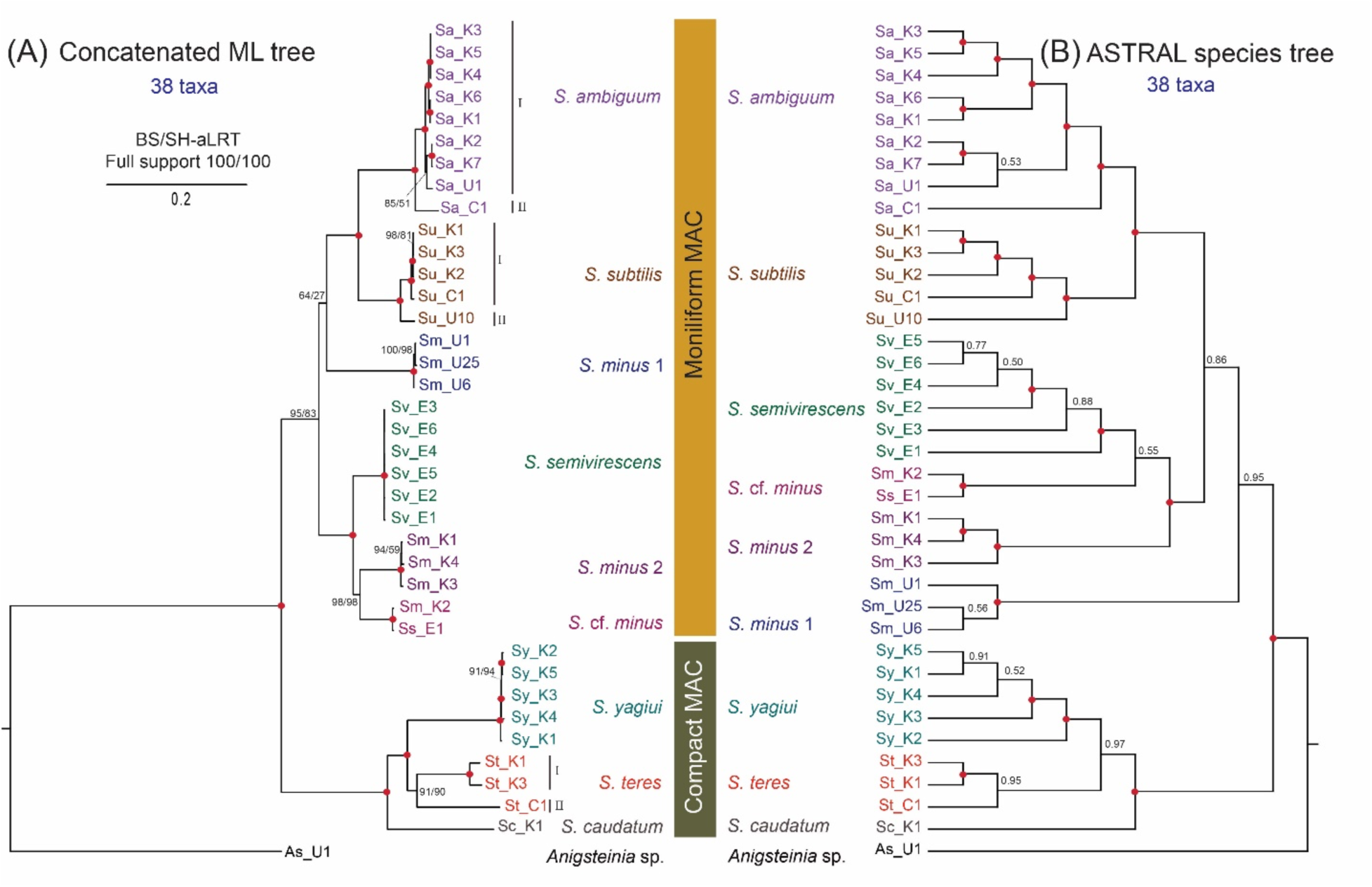
Phylogenomic relationships of *Spirostomum* species from 38 taxa and 265 protein-coding genes. (A) Maximum likelihood (ML) tree topology of a phylogenomic matrix comprising 62,043 unambiguously aligned amino acid residues was inferred using IQ-TREE under the LG +C60 + F + G4 model. (B) ASTRAL species tree topology inferred from 265 gene trees. One *Anigsteinia* sp. was used as an outgroup. Numbers at nodes are ML bootstrap values and SH-aLRT from 1,000 replicates (IQ-TREE) and local posterior probability (LPP) for ASTRAL are given on the nodes. Red solid circle on branches means maximal support. The scale bar corresponds to the number of nucleotide substitutions. (C= China; E = Europe; K = South Korea; U = USA).

The concatenated tree fully supports a moniliform clade of *Spirostomum*, with four taxa (*S. ambiguum, S. minus, S. semivirescens, S. subtilis*) (Fig. 3a). *Spirostomum ambiguum* and *S. subtilis* populations formed a monophyletic clade with full support. However, genetically divergent individuals from both *S. ambiguum* and *S. subtilis* clade were identified and our phylogenomic tree suggests the existence of cryptic species in both lineages. *S. minus* formed two well-supported clades: the *S. minus* clade 1 received a high support and falls sister to *S. ambiguum* + *S. subtils* clade with low support (64% UFBoot and 27% SH-aLTR), whereas the *S. minus* 2 clade formed a monophyletic clade with *Spirostomum* cf. *minus* with high statistical support (98% UFBoot and 98% SH-aLTR), and clustering together with *S. semivirescens* with full support. In the second major clade (compact macronucleus), *S. yagiui* populations were monophyletic and well supported. Individuals assigned to *S. teres* formed two distinct intraspecific lineages and *S. teres* (St_C1, isolates from China) with two other *S. teres* (St_K1 and St_K3, isolates from South Korea) formed a clade that obtained moderate statistical support (91% UFBoot and 90% SH-aLTR). *Sprirostomum caudatum* is the deepest branching taxa in the second major clade and formed a fully supported sister relationship with a clade comprising *S. yagiui* and *S. teres*.

Based on the concatenated 265 protein-coding genes dataset, the neighbor-net split (Fig. 4) indicated the existence of non-vertical signals that may be caused by deep incomplete lineage sorting, recombination, or hybridization, convergent evolution. Specifically, two conflicting noisy splits were identified. One was observed in the *S. minus*1 clade, which was linked with *S. minus* 2 + *Spirostomum* cf. *minus* and *Anigsteinia* through small net-like splits. The other split was in the *S. teres* clade, where two *teres* lineages were connected with *S. yagiui* population in a mesh-like manner made the relationships even more complicated.

**Figure 4.**
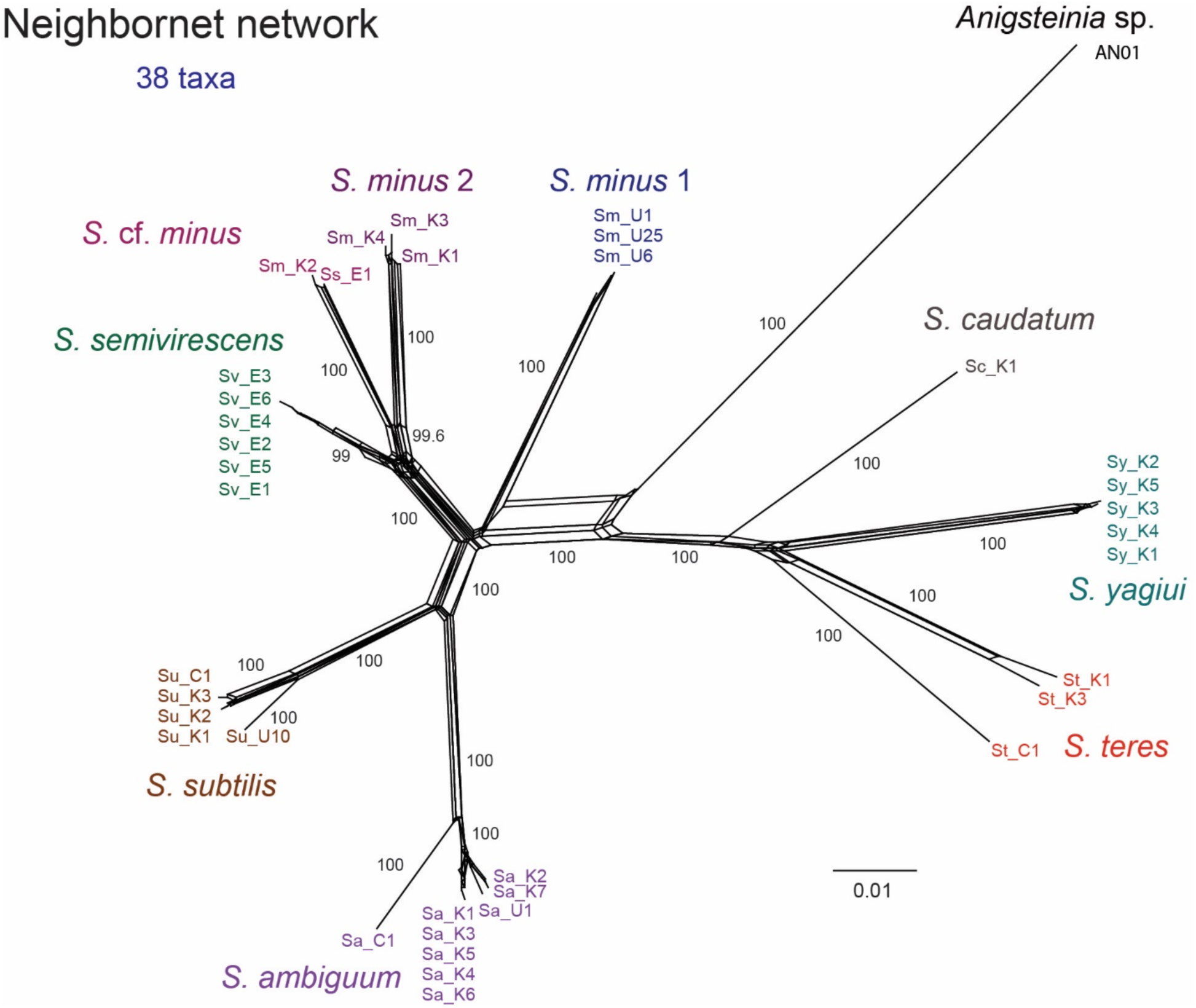
Phylogenetic network inferred from the concatenated alignment of 265 protein-coding genes using the neighbor net algorithm and uncorrected distances. Numbers along edges are bootstrap support values coming from 1,000 replicates. Values < 50% are not shown.

### Coalescent-based species tree using ASTRAL

To estimate the species tree using ASTRAL, we used the same number of genes as the concatenated IQ-TREE. Using ASTRAL, we estimated a species tree from the 265 gene trees (Fig. 3b). The resulting coalescent tree revealed similar relationships as the concatenated tree, and the discordant results received poorly supported nodes with the exception, in the ASTRAL species tree, *S. minus* 1 clade falls sister to the other early-branching taxa of moniliform MAC clade and received higher local posterior probability (LPP = 0.95) (Fig. 3b). Notably, *Spirostomum* cf. *minus* was more closely related to *S. semivirescens*, with poor statistical support (LPP = 0.55).

### Coalescent based species delimitation and species tree analysis using BPP

To determine whether the genetically divergent lineages found in IQ-TREE and ASTRAL analyses represent distinct species, we utilized the MSC model in BPP. Two different datasets (100 and 265 loci) were used for species delimitation analysis using A11 algorithm (Table 2, Supplementary Table 8, 9) We conducted the analysis multiple times and obtained consistent results, indicating that the chains were mixed properly. This ensures the reliability of the results (Yang 2015; Flouri et al. 2018). The species delimitation approach (A11) using BPP supports a species delimitation scheme where 13 lineages represent (including outgroup taxa) distinct species with full statistical support (posterior probability =1.00) in both 100 loci and 265 loci datasets (Fig. 5a). Further, we treated these all *Spirostomum* populations as 12 distinct lineages and used for species tree analysis using BPP.

**Figure 5.**
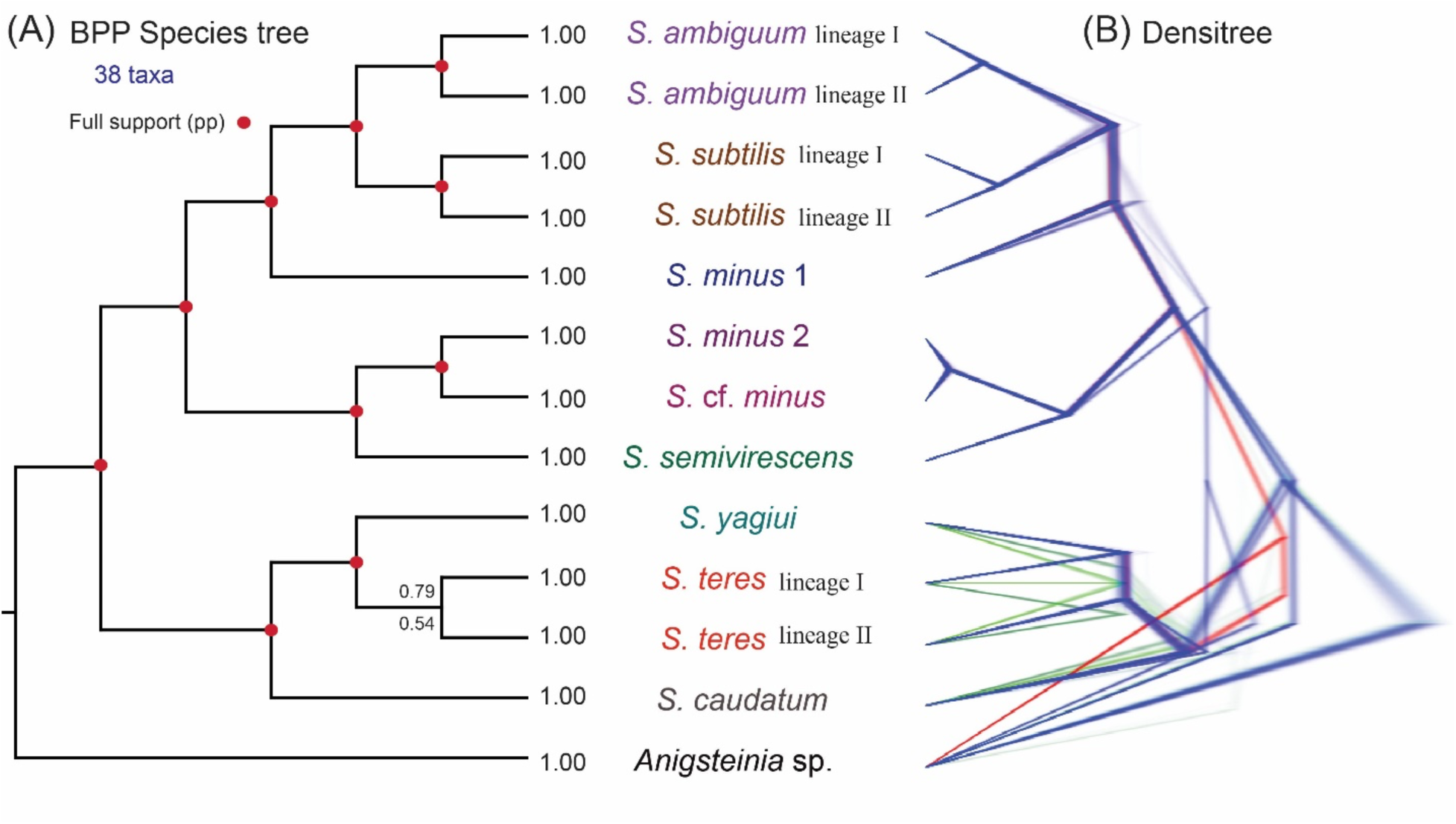
Multispecies coalescent-based species trees inferred using BPP. (A) Species tree using A01 approach with 265 loci datasets and 100 loci of 38 taxa assigned into 13 lineages. Posterior probability (PP) values are given at each node, upper value at the node indicates the PP of 265 loci and lower value at the node indicates the PP of 100 loci. Fully supported branches are marked with solid red circles. (B) DensiTree plot for 200K MCMC trees of 265 loci. Blue, red, and green colors represent the first, second, and third most common topologies, respectively, whereas gray represents other topologies.

**Table 2.** List of analyses performed in the present study.

| Analysis | No. of taxa | No. of genes | Description |
| --- | --- | --- | --- |
| 18S-ITS-28S rRNA tree | 136 | 3 | 130 *Spirostomum* + 1 *Pseudoblepharisma* + 5 *Anigsteinia* populations |
| Concatenated tree | 38 | 265 | Concatenated data matrix of 265 protein-coding genes. 37 *Spirostomum* + 1 *Anigsteinia* populations |
| Neighornet network | 38 | 265 | Concatenated data matrix of 265 protein-coding genes. 37 *Spirostomum* + 1 *Anigsteinia* populations |
| ASTRAL species tree | 38 | 265 | Gene trees of 265 protein-coding genes were used to estimate the species tree under the MSC model. 37 *Spirostomum* + 1 *Anigsteinia* populations |
| BPP 265 loci | 38 | 265 | Multiple Sequence Alignments (MSA) of 265 protein-coding genes were used to estimate the species tree and species delimitation under the MSC model. 37 *Spirostomum* + 1 *Anigsteinia* populations |
| BPP 100 loci | 38 | 100 | Multiple Sequence Alignments (MSA) of 100 protein-coding genes were used to estimate the species tree and species delimitation under the MSC model. 37 *Spirostomum* + 1 *Anigsteinia* populations |

The species tree inferred from the 100 and 265 loci using BPP (A01 analysis) are generally concordant with the concatenated alignment tree (Figs. 3a, 5a). Based on the MSC approach, the tree topology received full statistical support for all nodes, except the relationships between two *S. teres* lineages receive poor statistical support in both BPP species trees (100 and 265 loci dataset; Table 2). The 265 loci received higher support than 100 loci (posterior probability 0.79 vs 0.54), which might be the data size influenced the node support.

The DensiTree (Fig. 5b) of all 200,000 MCMC trees from 265 loci showed conflicting relationships between *S. teres* species and *S. yagiui*. The results showed that the *S. teres* lineages (I and II) are closely related and formed a monophyletic relationship, as shown in the blue-colored majority gene tree. On the other hand, the green-colored gene trees suggested that the *S. teres* lineages also formed a close relationship with *S. yagiui*, which may be due to high incomplete lineage sorting during rapid speciation events and inducing incongruence between gene trees and species trees.

## Discussion

In this study, we used single-cell transcriptomes to elucidate the phylogeny and delimitate species in *Spirostomum*, a genus of single-celled eukaryotes whose careful study can illuminate the origin of an unusual purple-green photosymbiosis, as well as the basis of unique cellular traits such as ultrafast contractility and collective intercellular communication (Mathijssen et al. 2019; Muñoz-Gómez et al. 2021; Muñoz-Gómez and Hess 2023). Members of genus *Spirostomum* are highly contractile and not always adaptable to laboratory conditions, posing challenges for taxonomists to conduct detailed morphological studies. So far, only a few *Spirostomum* species have been described with detailed morphological descriptions and 18S rRNA gene sequence information, while other species have been reported with only morphological data or 18S rRNA gene sequence information (Boscaro et al. 2014). Therefore, the evolution and species delimitation of *Spirostomum* species remains questionable due to limited data availability. To unravel the evolutionary history of *Spirostomum*, we used super matrix-based phylogenetic inference to estimate a species tree with IQ-TREE and a species network with SplitsTree. We thus subsequently used the MSC model that accounts for incomplete lineage sorting to estimate a species tree and delimit species boundaries using ASTRAL and BPP. BPP has been used for species tree and species delimitation analyses using large genomic data of multicellular organisms but has not yet been tested in unicellular eukaryotes (Pie et al. 2019; Ramirez-Reyes et al. 2020; Mcguire et al. 2023). Our analyses of transcriptome data provide insights into the phylogeny of *Spirostomum* and define the boundaries for its constituent species.

### Species delimitation using transcriptomic data analyses

Using the MSC-based analyses, we identified 12 putative species in the genus *Spirostomum* (Fig. 5). *Spirostomum minus* should be divided at least into two distinct species, which contrasts with the poor support for this clade in previous analyses relied on rRNA gene sequences (Fernandes and Silva Neto 2013; Boscaro et al. 2014; Shazib et al. 2016, 2019; Chi et al. 2020, 2022). Conserved rRNA gene regions could lead to ambiguous results in taxonomic studies, and no reliable morphological key characters were identified to differentiate *S. minus* clade species. Our MSC-based species delimitation approach using BPP confirms three species within *S. minus,* and we do not know which one is the originally described *S. minus* species as data on type specimens are lacking. Therefore, molecular data from type species is required to define the valid *S. minus* species, as also suggested in our previous work (Shazib et al. 2019).

Coalescent-based species delimitation analysis supports the existence of two genetically distinct species within *S. teres*; a taxon found in fresh and brackish water environments and characterized by a compact ellipsoidal macronucleus (Boscaro et al. 2014). Only *S. teres* and *S. caudatum* have an ellipsoidal macronucleus but these species can be easily distinguished from each other by their body shape (posterior end of the body blunt vs. posterior end of the body slender). In previous studies, multiple *S. teres* individuals appeared as non-monophyletic in rRNA gene trees and split into several clades (Boscaro et al. 2014; Chi et al. 2020, 2020; Fernandes and Silva Neto 2013; Shazib et al. 2019, 2016) (Supplementary Fig. S1). Even the ITS2 secondary structure could not differentiate *S. teres* populations due to insufficient information in the rRNA gene sequences (Shazib et al. 2016). In our study, we only have three *S. teres* populations with rRNA gene and transcriptomic data available (Supplementary Table S2). These three populations fall into two clades in the rRNA tree but recovered as monophyletic taxa both in the 265 protein-coding gene concatenated tree and the BPP species tree (Figs. 2, 3, 5, Supplementary Fig. S2). However, species delimitation approaches (A11analysis) define *S. teres* as two distinct lineages. Like *S. minus*, knowing which population corresponds to the “true’’ *S. teres* is also essential. Therefore, sequence data from type species is required.

Our species delimitation analysis also indicates *S. ambiguum* and *S. subtilis* contain two distinct lineages (Fig. 5). However, we identified the presence of cryptic species within *S. ambiguum* populations based on CO1 gene in our previous study (11.63% sequence divergence among *ambiguum* populations), but CO1 gene was not ambiguously able to define cryptic species in *S. subtilis* (Shazib et al. 2019).

### Phylogenetic relationships among Spirostomum species

Analyses of transcriptome data enable the reconstruction of a well-resolved phylogeny of the genus *Spirostomum*. In our previous study, we attempted to improve the phylogeny of *Spirostomum* species by increasing the taxon sampling and using five genetic markers from the nuclear and mitochondrial genome (Shazib et al. 2019). This was a significant improvement compared to previous studies that relied solely on rRNA gene-based phylogeny. Nonetheless, some relationships in the phylogenetic tree remained unclear. Our current analyses based on the phylogenomic approach enhance our understanding of the evolutionary pattern of *Spirostomum* species. For example, phylogenomic approach supports the deep division of *Spirostomum* into two major clades: Major clade 1 includes the members with moniliform macronucleus (*S. ambiguum*, *S. minus*, *S*. cf. *minus*, *S. semivirescens*, *S. subtilis*), while major clade 2 members (*S. caudatum*, *S. dharwarensis*, *S. teres*, *S. yagiui*) possess compact (ellipsoidal or elongated) macronucleus (Figs. 3, 5) and supports the two major clade species hypotheses (Boscaro et al. 2014; Shazib et al. 2016, 2019; Chi et al. 2020).

The macronuclear pattern within the class Heterotrichea is variable, and the evolution of macronuclear pattern is complex at the genus level (Schmidt et al. 2007; Thamm et al. 2010; Taher et al. 2020). According to our analysis of the rRNA gene tree (Fig. 2), *Pseudoblepharisma tenue* is closely related to the *Spirostomum* genus, which is consistent with previous studies using 18S rRNA and multigene (rRNA genes + CO1) tree analyses (Muñoz-Gómez et al. 2021; Hines et al. 2022). In terms of morphology, *Pseudoblepharisma tenue* shares similarities with *Spirostomum* ciliates, as the cell becomes spiral and shows contractility under stress and can be misidentified as a *Spirostomum* species because of cell size and ellipsoidal macronuclear shape (close to *S. teres*) (Muñoz-Gómez et al. 2021). Currently, only two species of *Pseudoblepharisma* have been documented, both of which possess a ellipsoidal macronucleus (Muñoz-Gómez et al. 2021; Hines et al. 2022). We hypothesize that a compact macronucleus is an ancestral state, whereas a moniliform macronucleus might be derived, as observed in other heterotrich genera (*Blepharisma, Stentor*) (Schmidt et al. 2007; Thamm et al. 2010; Taher et al. 2020).

Phylogenomic analyses using both concatenation and coalescent approach suggest the presence of high ancestral genetic polymorphism within the first major clade (species with moniliform macronucleus) (Figs. 3-5). In contrast with previous hypotheses, phylogenomic data indicate that *S. minus* 2 clade is more closely related to *S. semivirescens* than *S. minus* 1 clade, also consistently depicting *S. ambiguum* and *S. subtilis* as sister taxa (Figs. 3a, 4, 5). In previous molecular studies, *S. minus* appeared to be two morphologically cryptic species and their monophyly and/or separation (*minus* 1 and 2 clades) received poor support based on the rRNA gene phylogeny (Shazib et al. 2019), but the coalescent-based approach received more moderate support when *S. minus* splits in two different clades (Fig. 4 in Shazib et al. 2019). Phylogenetic analyses based on five genetic markers were limited to *S. minus* 2 clade; no data were available for *S. minus* 1 clade species or *S. semivirescens*. In our phylogenomic analyses, *Spirostomum* cf. *minus* like species is closely related to *S. minus* 2 clade species. Both concatenated analysis and coalesced based species tree support their monophyletic relationships (Figs. 3a, 5). However, coalescent-based species delimitation analysis suggests that *S. minus* 2 clade and *Spirostomum* cf. *minus* are distinct species. The evolutionary relationship within the clade with a moniliform macronucleus seems to be complex, and our study confirms that, minus-like character seems to have evolved independently at least two times in a convergent manner.

In the present study, concatenation and coalescent-based analyses indicate that *S. caudatum* is an early branching lineage within the second clade of *Spirostomum* (with ellipsoidal macronuclei; Figs. 3, 4, 5). *Spirostomum yagiui* is a monophyletic taxon and forms a sister relationship with *S. teres*. In our phylogenomic analyses, the populations under the name of *S. teres* recovered as monophyletic taxa but were non-monophyly in the rRNA gene tree and split in several clades (Figs. 2-5, Supplementary Fig. S2). Our result is congruent with the result from Shazib et al. (2019) and supports the hypothesis of Boscaro et al. (2014) that *S. yagiui* and *S. dharwarenis* (with an elongated macronucleus) originated during the rapid of a *S. teres*-like ancestor with an ellipsoidal macronucleus. Overall, our current data suggest that *S. teres* represent two different lineages. Transcriptomic or genomic data from other *S. teres* populations are needed to verify the monophyly of *S. teres* at a phylogenomic level.

## Conclusion

Our phylogenomic species tree and species delimitation approach, using transcriptomic data, allowed us to resolve the evolutionary relationships between *Spirostomum* species robustly. Our results confirm that *Spirostomum* species with a moniliform and compact macronucleus form distinct lineages, which is consistent with previous studies (Boscaro et al. 2014; Shazib et al. 2016, 2019). We furthermore tested the MSC model implemented in BPP on scRNA-seq data from populations representing seven *Spirostomum* morphospecies for the first time. This confirmed three distinct *S. minus* like species and *S. teres* represent two different lineages. Further revealed cryptic species in *S. ambiguum* and *S. subtilis*, which had not been previously suggested by rRNA gene trees. Our exploratory network analyses also suggest the occurrence of incomplete lineage sorting in the evolutionary history of the genus *Spirostomum*. The MSC-based and concatenation-based tree agreed with each other and suggest that BPP is a valuable tool for species tree inference and delimitation in single-celled eukaryotes. Overall, our fine sampling of closely related *Spirostomum* populations and wide scRNA-seq allowed us to demonstrate the hidden crypticity of species within the genus *Spirostomum*, and to resolve and provide much stronger support than hitherto to the phylogeny of this important ciliate genus.

## Data availability

Raw transcriptome reads are deposited in GenBank under the BioProject number PRJNA1113686, along with the rRNA gene sequences of species (XXXXXX–XXXXXX). The workflow to reconstruct rDNA from transcriptomic data is available at https://github.com/shahed30/SeqToDNA.

## Acknowledgments

We thank Ying Yan and Xyrus Maurer-Alcalá for collecting ciliates, and Angela Jiang, and Emma Schumacher for helping in writing additional scripts for the pipeline in Katz’s laboratory, Smith College. This work was supported by the grants from the NRF Korea funded by the Ministries of Science and ICT, and Education of the Republic of Korea under Grants 2017R1D1A2B03032963 (awarded to S.U.A.S) and 2018R1A2B6007973, 2021R1I1A2048744 (awarded to M.K.S). L.A.K is supported by grants from the NIH (R15HG010409) and NSF (OCE-1924570).

## Supplementary materials

**Supplementary Figure S1**: Six morphospecies of the genus *Spirostomum* isolated from different habitats in South Korea. Morphology of living cells (A-C) Species with moniliform macronucleus, and (D-F) species with compact macronucleus. (A) *Spirostomum ambiguum* has a moniliform macronucleus, the peristome occupying about 50% of the body length, and the posterior body end is truncated. (B) *S. subtilis* has a moniliform macronucleus, the peristome occupies about 35% of the body length, and the posterior body end is truncated. (C) *S. minus* has a moniliform macronucleus, the peristome occupying about 40% of the body length, and the posterior body end is truncated (D) *S. yagiui* has an elongated macronucleus, the peristome occupying about 42% of the body length, and the posterior body end is truncated. (E) *S. teres* has an ellipsoid macronucleus, the peristome occupies about 38% of the body length, and the posterior body end is truncated. (F) *S. caudatum* has an ellipsoid macronucleus, the peristome occupying about 35% of the body length, and a tail-like posterior body end. MAC, macronucleus; CV, contractile vacuole; CY, cytostome. Scale bars: 500 µm (A, B), 100 µm (C, D, F), 50 m (E).

**Supplementary Figure S2.** Maximum likelihood (ML) analysis using IQ-TREE of genus *Spirostomum*. The topology is based on 136 18S-ITS-28S rRNA gene sequences and 2,459 bp data matrix using the TIM2e+I+G4 model. Numbers at nodes are ML bootstrap values and SH-aLRT from 1,000 replicates. The scale bar corresponds to the number of nucleotide substitutions.

**Supplementary Figure S3.** Reconstruction of rDNA from Su_K1 transcriptome data using different read mapping software. PCR Sanger sequencing of Su_K1 used as a reference sequence for read mapping and contig assembling. BWA read mappers generated small and highly diverged contigs and were not included in this comparison analysis. Only contigs from BBMap, Bowtie 2, Hisat2 and q40 map quality were used here. Nucleotide different from the reference sequence (Su_K1 Sanger sequencing contig) were highlighted with red box.

**Supplementary Table S1.** List of primers used for PCR amplification and sequencing of rRNA gene.

**Supplementary Table S2**. Dataset and sequence information of 139 Spirostomidae ciliates used for rRNA and phylogenomic analyses (RM, contig reconstructed using read mapping).

**Supplementary Table S3.** BUSCO genome completeness analysis of 38 Spirostomidae ciliates used in this study.

**Supplementary Table S4.** rDNA reconstruction from WTA data using different mapping algorithm. For the comparisons here, we used *Spirostomum subtilis* (Su_K1) as a reference.

**Supplementary Table S5.** Characteristics of rRNA gene sequences of 38 Spirostomidae populations used in this study (RM, contig reconstructed using read mapping).

**Supplementary Table S6.** Reconstructed rRNA gene sequences using read mapping approach and reference sequences.

**Supplimentary Table S7.** Species delimitation (A11) and species tree (A01) analyses were conducted using different prior settings on 100 loci and 265 loci datasets to determine the best prior settings fit the dataset.

**Supplementary Table S8**. List of 265 protein-coding genes and species distribution across gene families (GFs). These 265 genes were utilized for phylogenomic and BPP analyses. Gene Family defined by OrthoMCL number followed by subclade number (e.g., OG5_126891_1, where 126891 is the ortholog group, and subclade number 1 contains the maximum number of target taxa).

**Supplementary Table S9**. List of 100 protein-coding genes and species distribution across gene families (GFs). These 100 genes were utilized for species tree analyses and species delimitation using BPP. Gene Family defined by OrthoMCL number followed by subclade number (e.g., OG5_126891_1, where 126891 is the ortholog group, and subclade number 1 contains the maximum number of target taxa).

## References

Akter S., Shazib S.U.A., Shin M.K. 2020. *Segnochrobactrum spirostomi* gen. nov., sp. nov., isolated from the ciliate *Spirostomum yagiui* and description of a novel family, *Segnochrobactraceae* fam. nov. within the order *Rhizobiales* of the class *Alphaproteobacteria*. Int. J. Syst. Evol. Microbiol. 70:1250–1258.

Andrews S. 2010. FastQC: A Quality Control Tool for High Throughput Sequence Data. Available online at: http://www.bioinformatics.babraham.ac.uk/projects/fastqc

Bankevich A., Nurk S., Antipov D., Gurevich A.A., Dvorkin M., Kulikov A.S., Lesin V.M., Nikolenko S.I., Pham S., Prjibelski A.D., Pyshkin A.V., Sirotkin A.V., Vyahhi N., Tesler G., Alekseyev M.A., Pevzner P.A. 2012. SPAdes: A New Genome Assembly Algorithm and its Applications to Single-Cell Sequencing. J. Comput. Biol. 19:455–477.

Berger H., Foissner W. 2003. Illustrated guide and ecological notes to ciliate indicator species (Protozoa, Ciliophora) in running waters, lakes, and sewage plants. In: Steinberg, C., Calmano, W., Klapper, H., Wilken, R.-D. (Eds.), Handbuch Angewandte Limnologie. Ecomed Verlag, Landsberg am Lech, 17. Erg.-Lieferung 10/03, p. 1–160.

Berger H., Foissner W., Kohmann F. 1997. Bestimmung und Ökologie der Mikrosaprobien nach DIN 38410. Gustav Fischer, Stuttgart, Jena, Lübeck, Ulm.

Boscaro V., Carducci D., Barbieri G., Senra M.V.X., Andreoli I., Erra F., Petroni G., Verni F., Fokin S.I. 2014. Focusing on Genera to Improve Species Identification: Revised Systematics of the Ciliate *Spirostomum*. Protist. 165:527–541.

Bouckaert R.R. 2010. DensiTree: making sense of sets of phylogenetic trees. Bioinformatics. 26:1372–1373.

Bradley M.W., Esteban G.F., Finlay B.J. 2010. Ciliates in chalk-stream habitats congregate in biodiversity hot spots. Res. Microbiol. 161:619–625.

Bryant D. 2003. Neighbor-Net: An Agglomerative Method for the Construction of Phylogenetic Networks. Mol. Biol. Evol. 21:255–265.

Bushmanova E., Antipov D., Lapidus A., Prjibelski A.D. 2019. rnaSPAdes: a *de novo* transcriptome assembler and its application to RNA-Seq data. Gigascience. 8:giz100.

Bushnell B. 2014. BBMap: A Fast, Accurate, Splice-Aware Aligner. Available at: https://sourceforge.net/projects/bbmap/.

Bushnell B., Rood J., Singer E. 2017. BBMerge – Accurate paired shotgun read merging via overlap. PLoS One. 12:e0185056.

Capella-Gutiérrez S., Silla-Martínez J.M., Gabaldón T. 2009. trimAl: a tool for automated alignment trimming in large-scale phylogenetic analyses. Bioinformatics. 25:1972–1973.

Cerón-Romero M.A., Maurer-Alcalá X.X., Grattepanche J.-D., Yan Y., Fonseca M.M., Katz L.A. 2019. PhyloToL: A Taxon/Gene-Rich Phylogenomic Pipeline to Explore Genome Evolution of Diverse Eukaryotes. Mol. Biol. Evol. 36:1831–1842.

Chen X., Kim J.H., Shazib S.U.A., Kwon C.B., Shin M.K. 2017. Morphology and molecular phylogeny of three heterotrichid species (Ciliophora, Heterotrichea), including a new species of Anigsteinia. Eur. J. Protistol. 61:278–293.

Chen X., Shazib S.U.A., Kim J.H., Kim M.S., Shin M.K. 2018. New contributions to *Gruberia lanceolata* (Gruber, 1884) Kahl, 1932 based on analyses of multiple populations and genes (Ciliophora, Heterotrichea, Gruberiidae). Eur. J. Protistol. 65:16–30.

Chi Y., Duan L., Luo X., Cheng T., Warren A., Huang J., Chen X. 2020. A new contribution to the taxonomy and molecular phylogeny of three, well-known freshwater species of the ciliate genus *Spirostomum* (Protozoa: Ciliophora: Heterotrichea). Zool. J. Linn. Soc. 189:158–177.

Chi Y., Wang Z., Ye T., Wang Y., Zhao J., Song W., Bourland W.A., Chen X. 2022. A new contribution to the taxonomy and phylogeny of the ciliate genus *Spirostomum* (Alveolata, Ciliophora, Heterotrichea), with comprehensive descriptions of two species from wetlands in China. Water Biol. Secur. 1:100031.

Clamp J.C., Lynn D.H. 2017. Investigating the biodiversity of ciliates in the ‘Age of Integration’. Eur. J. Protistol. 61:314–322.

Dopheide A., Lear G., Stott R., Lewis G. 2008. Molecular Characterization of Ciliate Diversity in Stream Biofilms. Appl. Environ. Microbiol. 74:1740–1747.

Esteban G.F., Bradley M.W., Finlay B.J. 2009. A case-building *Spirostomum* (Ciliophora, Heterotrichida) with zoochlorellae. Eur. J. Protistol. 45:156–158.

Fernandes N.M., Silva Neto I.D.D. 2013. Morphology and 18S rDNA gene sequence of *Spirostomum minus* and *Spirostomum teres* (Ciliophora: Heterotrichea) from Rio de Janeiro, Brazil. Zoologia (Curitiba). 30:72–79.

Finlay B.J. 2002. Global Dispersal of Free-Living Microbial Eukaryote Species. Science. 296:1061–1063.

Flouri T., Jiao X., Rannala B., Yang Z. 2018. Species Tree Inference with BPP Using Genomic Sequences and the Multispecies Coalescent. Mol. Biol. Evol. 35:2585–2593.

Flouri T., Jiao X., Rannala B., Yang Z. 2020a. A Bayesian Implementation of the Multispecies Coalescent Model with Introgression for Phylogenomic Analysis. Mol. Biol. Evol. 37:1211–1223.

Flouri T., Rannala B., Yang Z. 2020b. A Tutorial on the Use of BPP for Species Tree Estimation and Species Delimitation. In: Scornavacca C., Delsuc F., Galtier N., editors. Phylogenetics in the Genomic Era. No commercial publisher | Authors open access book. p. 5.6:1–5.6:16.

Foissner W., Berger H., Kohmann F. 1992. Taxonomische und ökologische Revision der Ciliaten des Saprobiesystems—Band II: Peritrichia, Heterotrichida, Odontostomatida. Informationsberichte des Bayer. Landesamtes für Wasserwirtschaft 5/92, 1–502.

Fokin S.I. 2012. Frequency and biodiversity of symbionts in representatives of the main classes of Ciliophora. Eur. J. Protistol. 48:138–148.

Fokin S.I., Schweikert M., Brümmer F., Görtz H.-D. 2005. *Spirostomum* spp. (Ciliophora, Protista), a suitable system for endocytobiosis research. Protoplasma. 225:93–102.

Guindon S., Dufayard J.-F., Lefort V., Anisimova M., Hordijk W., Gascuel O. 2010. New Algorithms and Methods to Estimate Maximum-Likelihood Phylogenies: Assessing the Performance of PhyML 3.0. Syst. Biol. 59:307–321.

Hakenkamp C.C., Morin A. 2000. The importance of meiofauna to lotic ecosystem functioning. Freshwater Biol. 44:165–175.

Hines H.N., McCarthy P.J., Esteban G.F. 2022. A Case Building Ciliate in the Genus *Pseudoblepharisma* Found in Subtropical Fresh Water. Diversity. 14:174.

Hines H.N., Onsbring H., Ettema T.J.G., Esteban G.F. 2018. Molecular Investigation of the Ciliate *Spirostomum semivirescens*, with First Transcriptome and New Geographical Records. Protist. 169:875–886.

Hoang D.T., Chernomor O., von Haeseler A., Minh B.Q., Vinh L.S. 2018. UFBoot2: Improving the Ultrafast Bootstrap Approximation. Mol. Biol. Evol. 35:518–522.

Huang J., Flouri T., Yang Z. 2020. A Simulation Study to Examine the Information Content in Phylogenomic Data Sets under the Multispecies Coalescent Model. Mol. Biol. Evol. 37:3211–3224.

Huson D.H. 1998. SplitsTree: analyzing and visualizing evolutionary data. Bioinformatics. 14:68–73.

Huson D.H., Bryant D. 2006. Application of Phylogenetic Networks in Evolutionary Studies. Mol. Biol. Evol. 23:254–267.

Jang S.W., Kwon C.-B., Shin M.K. 2012. First Records of Two *Spirostomum* Ciliates (Heterotrichea: Heterotrichida: Spirostomidae) from Korea. Animal Syst. Evol. Divers. 28:29–35.

Katoh K., Standley D.M. 2013. MAFFT Multiple Sequence Alignment Software Version 7: Improvements in Performance and Usability. Mol. Biol. Evol. 30:772–780.

Kearse M., Moir R., Wilson A., Stones-Havas S., Cheung M., Sturrock S., Buxton S., Cooper A., Markowitz S., Duran C., Thierer T., Ashton B., Meintjes P., Drummond A. 2012. Geneious Basic: An integrated and extendable desktop software platform for the organization and analysis of sequence data. Bioinformatics. 28:1647–1649.

Kim D., Paggi J.M., Park C., Bennett C., Salzberg S.L. 2019. Graph-based genome alignment and genotyping with HISAT2 and HISAT-genotype. Nat. Biotechnol. 37:907–915.

Kolisko M., Boscaro V., Burki F., Lynn D.H., Keeling P.J. 2014. Single-cell transcriptomics for microbial eukaryotes. Curr. Biol. 24:R1081–R1082.

Kreutz M., Foissner W. 2006. The Sphagnum Ponds of Simmelried in Germany : A biodiversity Hot-Spot for Microscopic Organisms. (Shaker Verlag GmbH, ed. 1, 2006).

Langmead B., Salzberg S.L. 2012. Fast gapped-read alignment with Bowtie 2. Nat. Methods. 9:357–359.

Li H., Durbin R. 2009. Fast and accurate short read alignment with Burrows–Wheeler transform. Bioinformatics. 25:1754–1760.

Liu Z., Hu S.K., Campbell V., Tatters A.O., Heidelberg K.B., Caron D.A. 2017. Single-cell transcriptomics of small microbial eukaryotes: limitations and potential. ISME J. 11:1282–1285.

Lynn D.H. 2008. The Ciliated Protozoa. Characterization, Classification, and Guide to the Literature. Springer, Dordrecht.

Manni M., Berkeley M.R., Seppey M., Zdobnov E.M. 2021. BUSCO: Assessing Genomic Data Quality and Beyond. Curr. Protocols. 1:e323.

Mathijssen A.J.T.M., Culver J., Bhamla M.S., Prakash M. 2019. Collective intercellular communication through ultra-fast hydrodynamic trigger waves. Nature. 571:560–564.

Mcguire J.A., Huang X., Reilly S.B., Iskandar D.T., Wang-Claypool C.Y., Werning S., Chong R.A., Lawalata S.Z.S., Stubbs A.L., Frederick J.H., Brown R.M., Evans B.J., Arifin U., Riyanto A., Hamidy A., Arida E., Koo M.S., Supriatna J., Andayani N., Hall R. 2023. Species Delimitation, Phylogenomics, and Biogeography of Sulawesi Flying Lizards: A Diversification History Complicated by Ancient Hybridization, Cryptic Species, and Arrested Speciation. Syst. Biol.:syad020.

Minh B.Q., Nguyen M.A.T., von Haeseler A. 2013. Ultrafast Approximation for Phylogenetic Bootstrap. Mol. Biol. .Evol. 30:1188–1195.

Minh B.Q., Schmidt H.A., Chernomor O., Schrempf D., Woodhams M.D., von Haeseler A., Lanfear R. 2020. IQ-TREE 2: New Models and Efficient Methods for Phylogenetic Inference in the Genomic Era. Mol. Biol. .Evol. 37:1530–1534.

Mirarab S., Reaz R., Bayzid Md.S., Zimmermann T., Swenson M.S., Warnow T. 2014. ASTRAL: genome-scale coalescent-based species tree estimation. Bioinformatics. 30:i541–i548.

Mukhtar I., Wu S., Wei S., Chen R., Cheng Y., Liang C., Chen J. 2020. Transcriptome Profiling Revealed Multiple rquA Genes in the Species of *Spirostomum* (Protozoa: Ciliophora: Heterotrichea). Front. Microbiol. 11:574285.

Muñoz-Gómez S.A., Hess S. 2023. Purple photosymbioses. Curr. Biol. 33:R167–R170.

Muñoz-Gómez S.A., Kreutz M., Hess S. 2021. A microbial eukaryote with a unique combination of purple bacteria and green algae as endosymbionts. Sci. Adv. 7:eabg4102.

Nałecz-Jawecki G., Sawicki J. 1999. Spirotox — A new tool for testing the toxicity of volatile compounds. Chemosphere. 38:3211–3218.

Nałęcz-Jawecki G., Wawryniuk M., Giebułtowicz J., Olkowski A., Drobniewska A. 2020. Influence of Selected Antidepressants on the Ciliated Protozoan *Spirostomum ambiguum*: Toxicity, Bioaccumulation, and Biotransformation Products. Molecules. 25:1476.

Onsbring H., Tice A.K., Barton B.T., Brown M.W., Ettema T.J.G. 2020. An efficient single-cell transcriptomics workflow for microbial eukaryotes benchmarked on *Giardia intestinalis* cells. BMC Genomics. 21:448.

Picelli S., Faridani O.R., Björklund Å.K., Winberg G., Sagasser S., Sandberg R. 2014. Full-length RNA-seq from single cells using Smart-seq2. Nat. Protoc. 9:171–181.

Pie M.R., Bornschein M.R., Ribeiro L.F., Faircloth B.C., McCormack J.E. 2019. Phylogenomic species delimitation in microendemic frogs of the Brazilian Atlantic Forest. Mol. Phylogenet. Evol. 141:106627.

Ramirez-Reyes T., Blair C., Flores-Villela O., Pinero D., Lathrop A., Murphy R. 2020. Phylogenomics and molecular species delimitation reveals great cryptic diversity of leaf-toed geckos (Phyllodactylidae: *Phyllodactylus*), ancient origins, and diversification in Mexico. Mol. Phylogenet. Evol. 150:106880.

Repak A., Isquith I. 1974. The systematics of the genus *Spirostomum* Ehrenberg, 1838. Acta. Protozool. 12:325–333.

Robert C.P., Casella G. 2004. Monte Carlo Statistical Methods. New York, NY: Springer New York.

Sayyari E., Mirarab S. 2016. Fast Coalescent-Based Computation of Local Branch Support from Quartet Frequencies. Mol. Biol. Evol. 33:1654–1668.

Schmidt S.L., Foissner W., Schlegel M., Bernhard D. 2007. Molecular Phylogeny of the Heterotrichea (Ciliophora, Postciliodesmatophora) Based on Small Subunit rRNA Gene Sequences. J. .Eukaryotic Microbiol. 54:358–363.

Schrallhammer M., Ferrantini F., Vannini C., Galati S., Schweikert M., Görtz H.-D., Verni F., Petroni G. 2013. ‘*Candidatus* Megaira polyxenophila’ gen. nov., sp. nov.: Considerations on Evolutionary History, Host Range and Shift of Early Divergent Rickettsiae. PLoS One. 8:e72581.

Shazib S.U.A., Vd’ačný P., Kim J.H., Jang S.W., Shin M.K. 2014. Phylogenetic relationships of the ciliate class Heterotrichea (Protista, Ciliophora, Postciliodesmatophora) inferred from multiple molecular markers and multifaceted analysis strategy. Mol. Phylogenet. Evol. 78:118–135.

Shazib S.U.A., Vďačný P., Kim J.H., Jang S.W., Shin M.K. 2016. Molecular phylogeny and species delimitation within the ciliate genus *Spirostomum* (Ciliophora, Postciliodesmatophora, Heterotrichea), using the internal transcribed spacer region. Mol. Phylogenet. Evol. 102:128–144.

Shazib S.U.A., Vďačný P., Slovák M., Gentekaki E., Shin M.K. 2019. Deciphering phylogenetic relationships and delimiting species boundaries using a Bayesian coalescent approach in protists: A case study of the ciliate genus *Spirostomum* (Ciliophora, Heterotrichea). Sci. Rep. 9:16360.

Shi C.-M., Yang Z. 2018. Coalescent-Based Analyses of Genomic Sequence Data Provide a Robust Resolution of Phylogenetic Relationships among Major Groups of Gibbons. Mol. Biol. Evol. 35:159–179.

Si Quang L., Gascuel O., Lartillot N. 2008. Empirical profile mixture models for phylogenetic reconstruction. Bioinformatics. 24:2317–2323.

Specht H. 1934. Aerobic respiration in *Spirostomum ambiguum* and the production of ammonia. J. Cell. Comp. Physiol. 5:319–333.

Taher M.A., Kabir A.S., Shazib S.U.A., Kim M.S., Shin M.K. 2020. Morphological Redescriptions and Molecular Phylogeny of Three *Stentor* Species (Ciliophora: Heterotrichea: Stentoridae) from Korea. Zootaxa. 4732:zootaxa.4732.3.6.

Thamm M., Schmidt S.L., Bernhard D. 2010. Insights into the Phylogeny of the Genus Stentor (Heterotrichea, Ciliophora) with Special Emphasis on the Evolution of the Macronucleus Based on SSU rDNA Data. Acta Protozool. 2010:149–157.

Thawornwattana Y., Seixas F.A., Yang Z., Mallet J. 2022. Full-Likelihood Genomic Analysis Clarifies a Complex History of Species Divergence and Introgression: The Example of the erato-sara Group of *Heliconius* Butterflies. Syst. Biol. 71:1159–1177.

Xu B., Yang Z. 2016. Challenges in Species Tree Estimation Under the Multispecies Coalescent Model. Genetics. 204:1353–1368.

Yang Z. 2015. The BPP program for species tree estimation and species delimitation. Curr. Zool. 61:854–865.

Yang Z., Rannala B. 2014. Unguided species delimitation using DNA sequence data from multiple Loci. Mol Biol Evol. 31:3125–3135.

Zhang C., Rabiee M., Sayyari E., Mirarab S. 2018. ASTRAL-III: polynomial time species tree reconstruction from partially resolved gene trees. BMC Bioinf. 19:153.

