## Supplementary material for "Phylogeny and species delimitation of ciliates in the genus *Spirostomum* (Class, Heterotrichea) using single-cell transcriptomes": Supplementary Figure S1.pdf

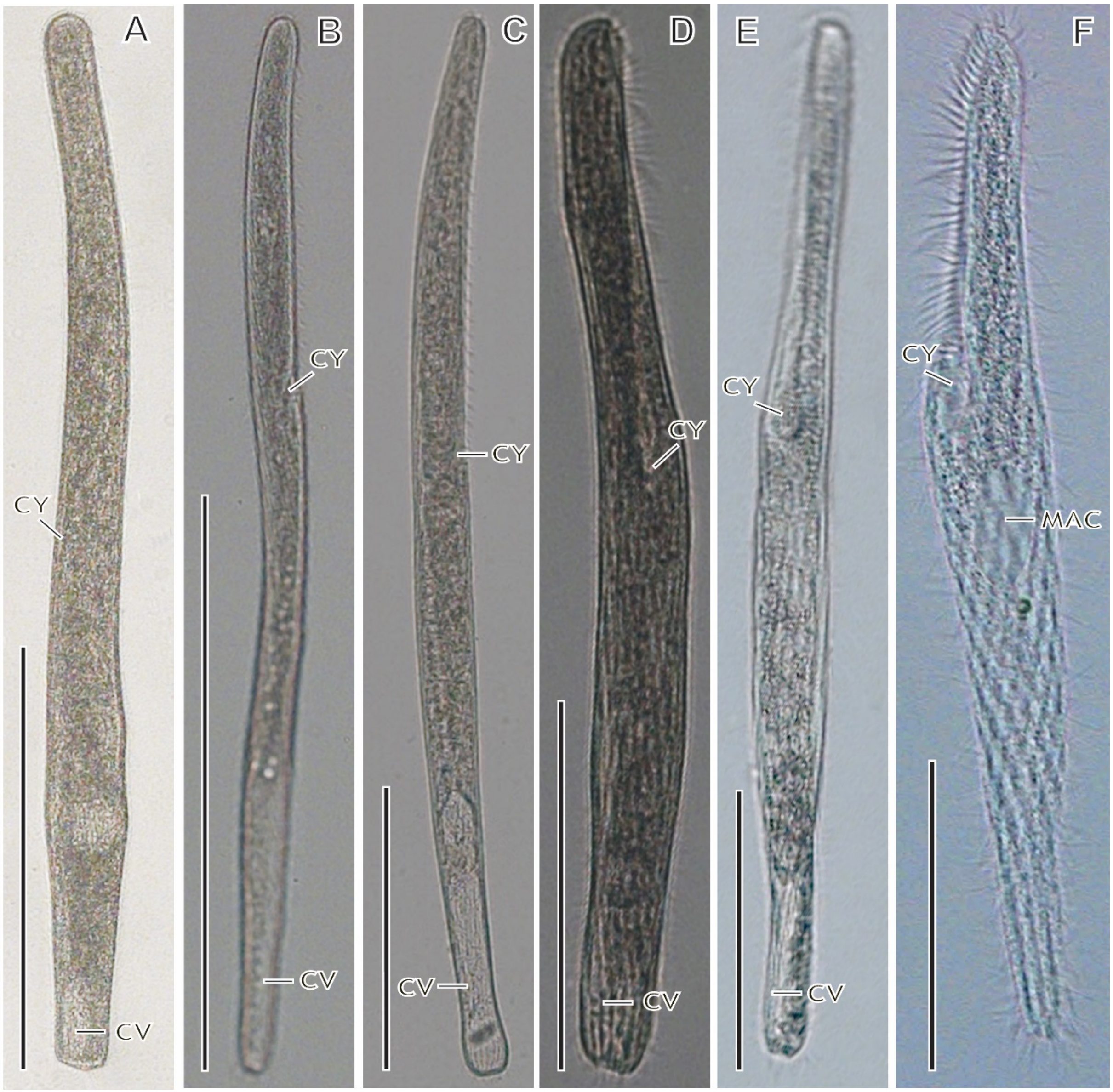


**Supplementary Figure S1**: Six morphospecies of the genus *Spirostomum* isolated from different habitats in South Korea. Morphology of living cells (A-C) Species with moniliform macronucleus, and (D-F) species with compact macronucleus. (A) *Spirostomum ambiguum* has a moniliform macronucleus, the peristome occupying about 50% of the body length, and the posterior body end is truncated. (B) *S. subtilis* has a moniliform macronucleus, the peristome occupies about 35% of the body length, and the posterior body end is truncated. (C) *S. minus* has a moniliform macronucleus, the peristome occupying about 40% of the body length, and the posterior body end is truncated (D) *S. yagiui* has an elongated macronucleus, the peristome occupying about 42% of the body length, and the posterior body end is truncated. (E) *S. teres* has an ellipsoid macronucleus, the peristome occupies about 38% of the body length, and the posterior body end is truncated. (F) *S. caudatum* has an ellipsoid macronucleus, the peristome occupying about 35% of the body length, and a tail-like posterior body end. MAC, macronucleus; CV, contractile vacuole; CY, cytostome. Scale bars: 500 µm (A, B), 100 µm (C, D, F), 50 m (E).

**References:**

Boscaro V., Carducci D., Barbieri G., Senra M.V.X., Andreoli I., Erra F., Petroni G., Verni F., Fokin S.I. 2014. Focusing on Genera to Improve Species Identification: Revised Systematics of the Ciliate *Spirostomum*. Protist. 165:527–541.

Chi Y., Duan L., Luo X., Cheng T., Warren A., Huang J., Chen X. 2020. A new contribution to the taxonomy and molecular phylogeny of three, well-known freshwater species of the ciliate genus *Spirostomum* (Protozoa: Ciliophora: Heterotrichea). Zool. J. Linn. Soc. 189:158–177.

Chi Y., Wang Z., Ye T., Wang Y., Zhao J., Song W., Bourland W.A., Chen X. 2022. A new contribution to the taxonomy and phylogeny of the ciliate genus *Spirostomum* (Alveolata, Ciliophora, Heterotrichea), with comprehensive descriptions of two species from wetlands in China. Water Biol. Secur. 1:100031.

Shazib S.U.A., Vďačný P., Kim J.H., Jang S.W., Shin M.K. 2016. Molecular phylogeny and species delimitation within the ciliate genus Spirostomum (Ciliophora, Postciliodesmatophora, Heterotrichea), using the internal transcribed spacer region. Mol. Phylogenet. Evol. 102:128–144.

Shazib S.U.A., Vďačný P., Slovák M., Gentekaki E., Shin M.K. 2019. Deciphering phylogenetic relationships and delimiting species boundaries using a Bayesian coalescent approach in protists: A case study of the ciliate genus *Spirostomum* (Ciliophora, Heterotrichea). Sci. Rep. 9:16360.
