## Supplementary material for "Phylogeny and species delimitation of ciliates in the genus *Spirostomum* (Class, Heterotrichea) using single-cell transcriptomes": Supplementary Figure S2.pdf

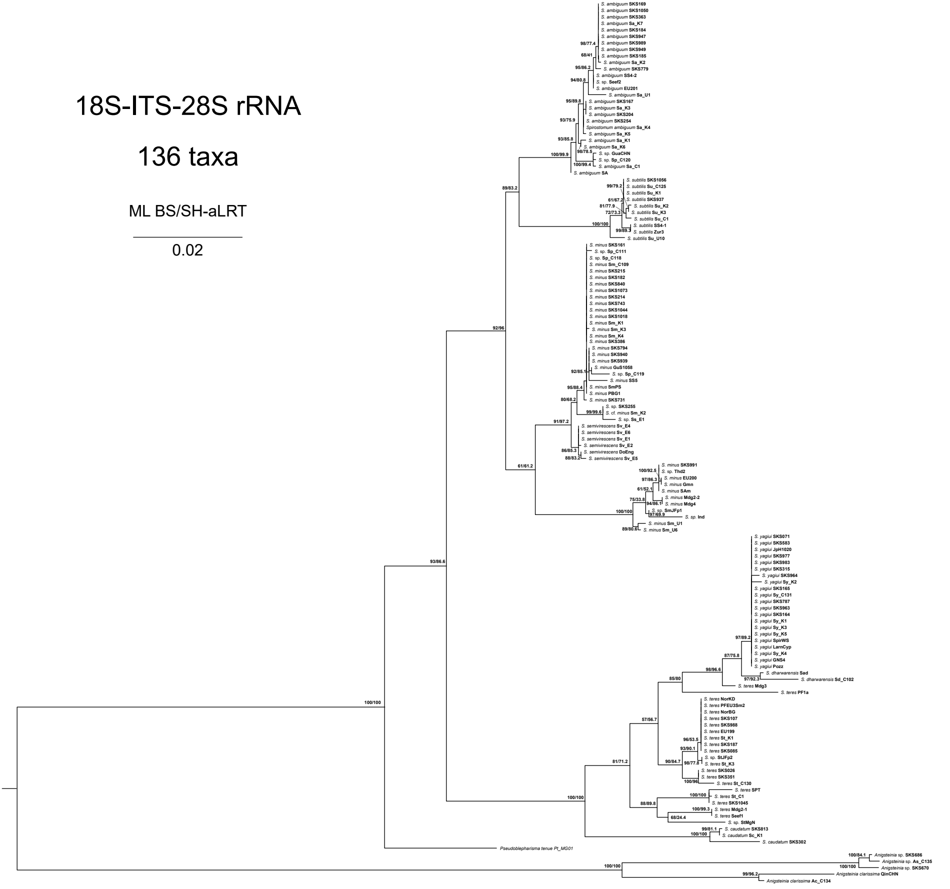


**Supplementary Figure S2**: Maximum likelihood (ML) analysis using IQ-TREE of genus *Spirostomum*. The topology is based on 136 18S-ITS-28S rRNA gene sequences and 2,459 bp data matrix using the TIM2e+I+G4 model. Numbers at nodes are ML bootstrap values and SH-aLRT from 1,000 replicates. The scale bar corresponds to the number of nucleotide substitutions.
