## Supplementary material for "Phylogeny and species delimitation of ciliates in the genus *Spirostomum* (Class, Heterotrichea) using single-cell transcriptomes": Supplementary Figure S3.pdf

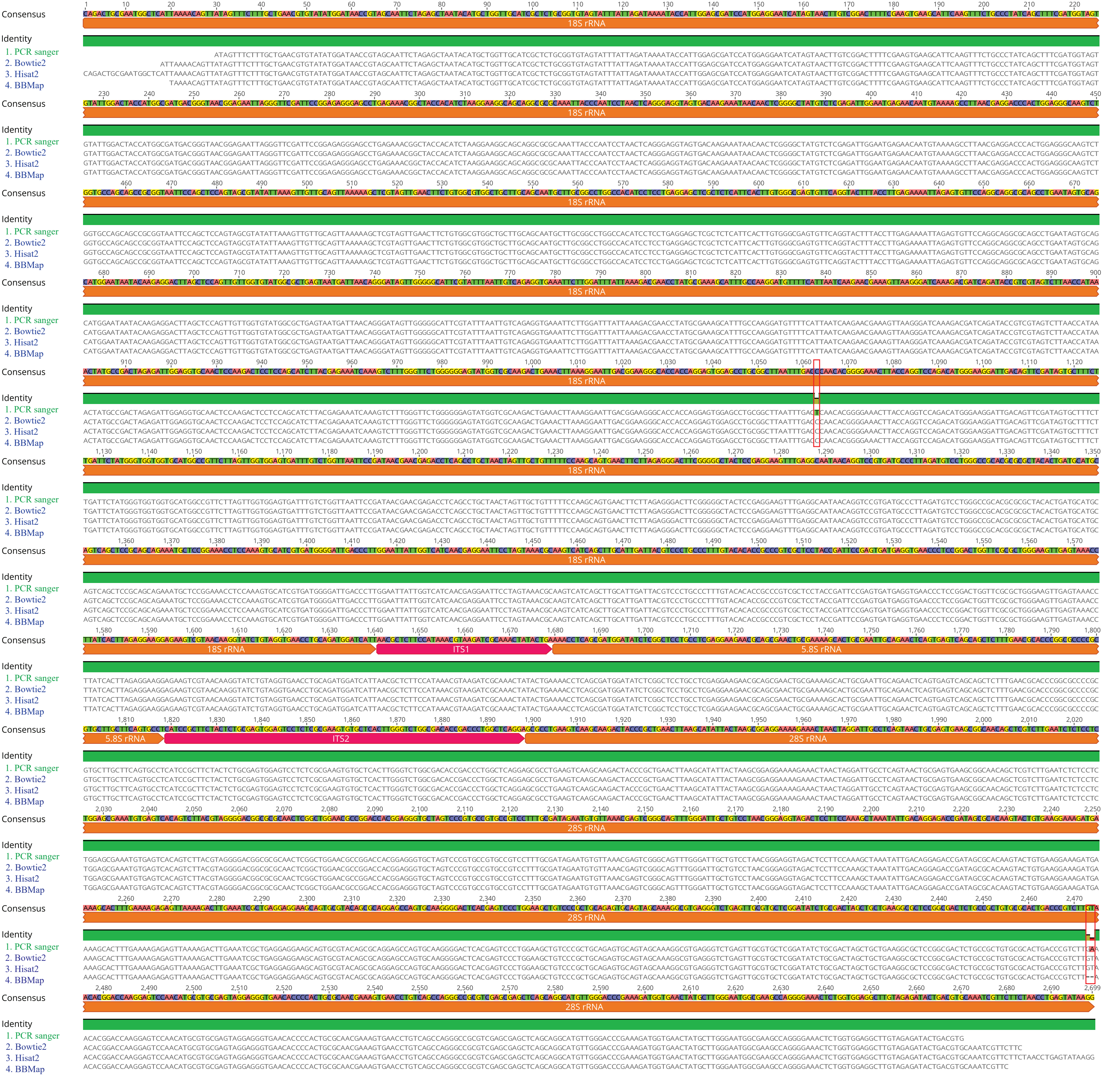


**Supplementary Figure S3:** Reconstruction of rDNA from Su_K1 transcriptome data using different read mapping software. PCR Sanger sequencing of Su_K1 used as a reference sequence for read mapping and contig assembling. BWA read mappers generated small and highly diverged contigs and were not included in this comparison analysis. Only contigs from BBMap, Bowtie 2, Hisat2 and q40 map quality were used here. Nucleotide different from reference sequences (Su_K1 Sanger sequencing contig) were highlighted with red box.
