## Supplementary material for "Phylogeny and species delimitation of ciliates in the genus *Spirostomum* (Class, Heterotrichea) using single-cell transcriptomes": Supplementary Table S1.docx

**Supplementary Table S1.** List of primers used for PCR amplification and sequencing of rRNA gene.

| **Primer Name** | **Sequence** | **Purpose** | **Reference** |
| --- | --- | --- | --- |
| Euk A | 5′ -AAC CTG GTT GAT CCT GCC AG-3′ | Primer for PCR and sequencing | (Medlin et al. 1988) |
| Euk B | 5′-CAC TTG GAC GTC TTC CTA GT-3′ | Primer for sequencing | (Medlin et al. 1988) |
| 18S 528F | 5′-CCG CGG TAA TTC CAG CTC-3′ | Primer for sequencing | (Alves-de-Souza et al. 2011) |
| ITS F | 5′-GTT CCC CTT GAA CGA GGA ATT C-3′ | Primer for sequencing | (Goggin and Murphy 2000) |
| ITS R | 5′-TAC TGA TAT GCT TAA GTT CAG CGG-3′ | Primer for sequencing | (Goggin and Murphy 2000) |
| D1D2 fwd1 | 5′-AGC GGG AGG AAA AGA AAC T-3′ | Primer for sequencing | (Sonnenberg et al. 2007) |
| D1D2 rev2 | 5′-ACG ATC GAT TTG CAC GTC AG-3′ | Primer for PCR and sequencing | (Sonnenberg et al. 2007) |

**References:**

Alves-de-Souza C., Cornet C., Nowaczyk A., Gasparini S., Skovgaard A., Guillou L. 2011. *Blastodinium* spp. infect copepods in the ultra-oligotrophic marine waters of the Mediterranean Sea. Biogeosciences. 8:2125–2136.

Goggin C., Murphy N. 2000. Conservation of sequence in the internal transcribed spacers and 5.8S ribosomal RNA among geographically separated isolates of parasitic scuticociliates (Ciliophora, Orchitophryidae). Dis. Aquat. Org. 40:79–83.

Medlin L., Elwood H.J., Stickel S., Sogin M.L. 1988. The characterization of enzymatically amplified eukaryotic 16S-like rRNA-coding regions. Gene. 71:491–499.

Sonnenberg R., Nolte A.W., Tautz D. 2007. An evaluation of LSU rDNA D1-D2 sequences for their use in species identification. Front. Zool. 4:6.
