## Supplementary material for "Phylogeny and species delimitation of ciliates in the genus *Spirostomum* (Class, Heterotrichea) using single-cell transcriptomes": Supplementary Table S4.docx

**Supplementary Table S4:** rDNA reconstruction from WTA data using different mapping softwares. For the comparisons here, we used *Spirostomum subtilis* (Su_K1) as a reference.

| **Mapper** |  |  |  | **Only mapped** |  |  |  | **q20** |  |  |  | **q40** |  |
| --- | --- | --- | --- | --- | --- | --- | --- | --- | --- | --- | --- | --- | --- |
|  |  |  | **Contig #** | **Reads #** | **Longest contig** |  | **Contig #** | **Reads #** | **Longest contig** |  | **Contig #** | **Reads #** | **Longest contig** |
| BBMap |  |  | 15 | 652,411 | 2,867 bp |  | 13 | 635,118 | 2,766 bp |  | 1 | 554,016 | 2,659 bp |
| BWA |  |  | 104 | 740,602 | 1,275 bp |  | 104 | 740,597 | 1,275 bp |  | 104 | 740,582 | 1,275 bp |
| Bowtie 2 |  |  | 3 | 578,248 | 2,689 bp |  | 2 | 566,246 | 2,688 bp |  | 1 | 528,264 | 2,666 bp |
| HISAT2 |  |  | 6 | 1,283,324 | 2,626 bp |  | 1 | 634,870 | 2,699 bp |  | 1 | 634,870 | 2,699 bp |
